# Structure Can Predict Function in the Human Brain: A Graph Neural Network Deep Learning Model of Functional Connectivity and Centrality Based on Structural Connectivity

**DOI:** 10.1101/2021.03.15.435531

**Authors:** Josh Neudorf, Shaylyn Kress, Ron Borowsky

## Abstract

Although functional connectivity and associated graph theory measures (e.g., centrality; how centrally important to the network a region is) are widely used in brain research, the full extent to which these functional measures are related to the underlying structural connectivity is not yet fully understood. Graph neural network deep learning methods have not yet been applied for this purpose, and offer an ideal model architecture for working with connectivity data given their ability to capture and maintain inherent network structure. This model applied here to predict functional connectivity and centrality from structural connectivity accounted for 89% of the variance in mean functional connectivity, 56% of the variance in individual-level functional connectivity, 99% of the variance in mean functional centrality, and 81% of the variance in individual-level functional centrality. This model provides a new benchmark for performance and represents a novel finding that functional centrality can be robustly predicted from structural connectivity. Regions of particular importance to the model’s performance as determined through lesioning are discussed, whereby regions with higher centrality have a higher impact on model performance. Future research on models of patient, demographic, or behavioural data can also benefit from this graph neural network method as it is ideally-suited for capturing connectivity and centrality in brain networks. These results have set a new benchmark for prediction of functional connectivity from structural connectivity, and models like this may ultimately lead to a way to predict functional connectivity in individuals who are unable to do fMRI tasks (e.g., non-responsive patients).

---

There is now widespread usage of functional connectivity and associated graph theory measures (e.g., centrality). Investigating to what extent the structural connectivity as measured by diffusion tensor imaging (DTI) can explain functional connectivity as measured by resting state functional magnetic resonance imaging (rsfMRI) is an important step towards understanding the structural basis of functional macroscale networks (the importance of this problem has recently been highlighted; Suárez et al., 2020). Some of the first steps towards understanding the relationship between structure and function showed moderate correspondence when examining direct structural connections, accounting for approximately 50% of the variance (Honey et al., 2009), or looking at a subset of connections (62%; Hagmann et al., 2008). However, when examining all functional connections, structural connectivity accounts for only 9% of the variance using linear regression by one account (Rosenthal et al., 2018). Novel graph theory measures calculated using structural connectivity have accounted for 23% of the variance in functional connectivity (Goñi et al., 2014), a combination of vector encodings of structural connectivity and deep learning with a fully connected network (FCN) have accounted for 36% of the variance in functional connectivity (Rosenthal et al., 2018), and simulated fMRI activation using a hybrid approach with both DTI and electroencephalography (EEG) data has accounted for 53% of the variance in functional connectivity (Schirner et al., 2018). Still, there remains a large amount of variance unaccounted for by these models if we are to determine the extent to which functional connectivity and measures of centrality based on functional connectivity offer insight into the properties of the underlying structural network. Recently, a feed-forward FCN deep learning model was able to demonstrate that mean structural connectivity as input could predict mean functional connectivity accounting for 81% of the variance, and that individual-level structural connectivity could predict individual-level functional connectivity accounting for 30% of the variance (Sarwar et al., 2021). These findings represent a large improvement in prediction of functional connectivity. Further deep learning models should be investigated to determine whether there is a converging upper limit on functional connectivity prediction, in order to provide a benchmark to work towards with more explicit mathematical and simulation methods.

There is a general consensus in the neuroscience community that rsfMRI functional connectivity measures represent the effective connectivity between regions in the brain. Effective connectivity describes the meaningful result of communication between regions that are sparsely connected by direct and indirect structural connections (white matter tracts). To support this view researchers have noted that patterns of connectivity seem to occur between regions that are expected to function together based on previous research and neuroanatomy (see van den Heuvel and Hulshoff Pol, 2010 for a review). Modern preprocessing methods are able to account for the effects of physiological factors (e.g., respiration and cardiac function), in order to increase the neuronal basis of the BOLD signal relative to noise (e.g., Birn et al., 2008; Chang et al., 2009; Falahpour et al., 2013; Golestani et al., 2015; Kassinopoulos and Mitsis, 2019; Salas et al., 2021). Even with these advances, coordinated functional BOLD signal (functional connectivity) has yet to be robustly linked to the organization of the underlying structural network. Searching for the upper limit in predicting functional connectivity from structure remains an important goal for investigating to what extent the rsfMRI functional connectivity is influenced by the connectivity of the structural architecture of the brain.

One issue with structural connectivity is that DTI tractography data is sparse, meaning the majority of values are zeros. On the other hand, functional connectivity data has many more non-zero values, even when thresholded. This difference occurs in part because many routes of communication between brain regions are indirect rather than direct, and is one reason functional connectivity has been widely used. Measures of indirect (‘effective’) connectivity are available using graph theory, including shortest path length, communicability (Estrada and Hatano, 2008), and novel measures designed for brain research (e.g., search information and path transitivity; Goñi et al., 2014), but research is still investigating the extent to which these measures are based on sound assumptions about how functional connectivity results from underlying structure at the macroscale.

Research employing graph theory measures have become an important focus of recent network neuroscience research involving structural and functional connectivity (see Fornito et al., 2013), including measures of centrality, which describe a region’s importance to the network. Some common centrality measures include degree centrality (number of connections to a region), eigenvector centrality (number of connections to a region weighted by the centrality of its neighbours), and PageRank centrality (a variant of eigenvector centrality developed for use in ranking web pages, with the advantage that it addresses the issue of eigenvector centrality sometimes being excessively high when a low degree node is connected to a high centrality node; Page et al., 1999). Figure 1A depicts an example graph with labels showing which node has the highest centrality for each of these measures. This example graph is relatively sparse (fewer connections) like the structural connectivity network of the brain, in comparison to functional connectivity networks, which are denser (highly connected). Functional connectivity centrality has been used to demonstrate age and sex related differences (Zuo et al., 2012), differences between patient and control groups (for patients with schizophrenia, Chen et al., 2015; bipolar disorder, Deng et al., 2019, Zhou et al., 2017; retinitus pigmentosa, Lin et al., 2021; and diabetic optic neuropathy, Xu et al., 2020), and differences related to genotype (Wink et al., 2018). Structural connectivity centrality has also been used to demonstrate differences between patient and control groups (for patients with prenatal alcohol exposure, Long et al., 2020; traumatic brain injury, Raizman et al., 2020; gut inflamation, Turkiewicz et al., 2021; and brain tumours, Yu et al., 2016), and to demonstrate a relationship between structural centrality and functional complexity (e.g., Hurst exponent; inversely related to fractal dimension, where a fractal dimension exceeding the topological dimension of the signal indicates complex functional activity) suggesting that regions integrating information from many sources have more complex functional activity (Neudorf et al., 2020). An important question that has not yet been explored, to our knowledge, is to what extent variance in functional connectivity-based centrality measures can be accounted for by structural connectivity and structural connectivity-based centrality.

**Figure 1.**
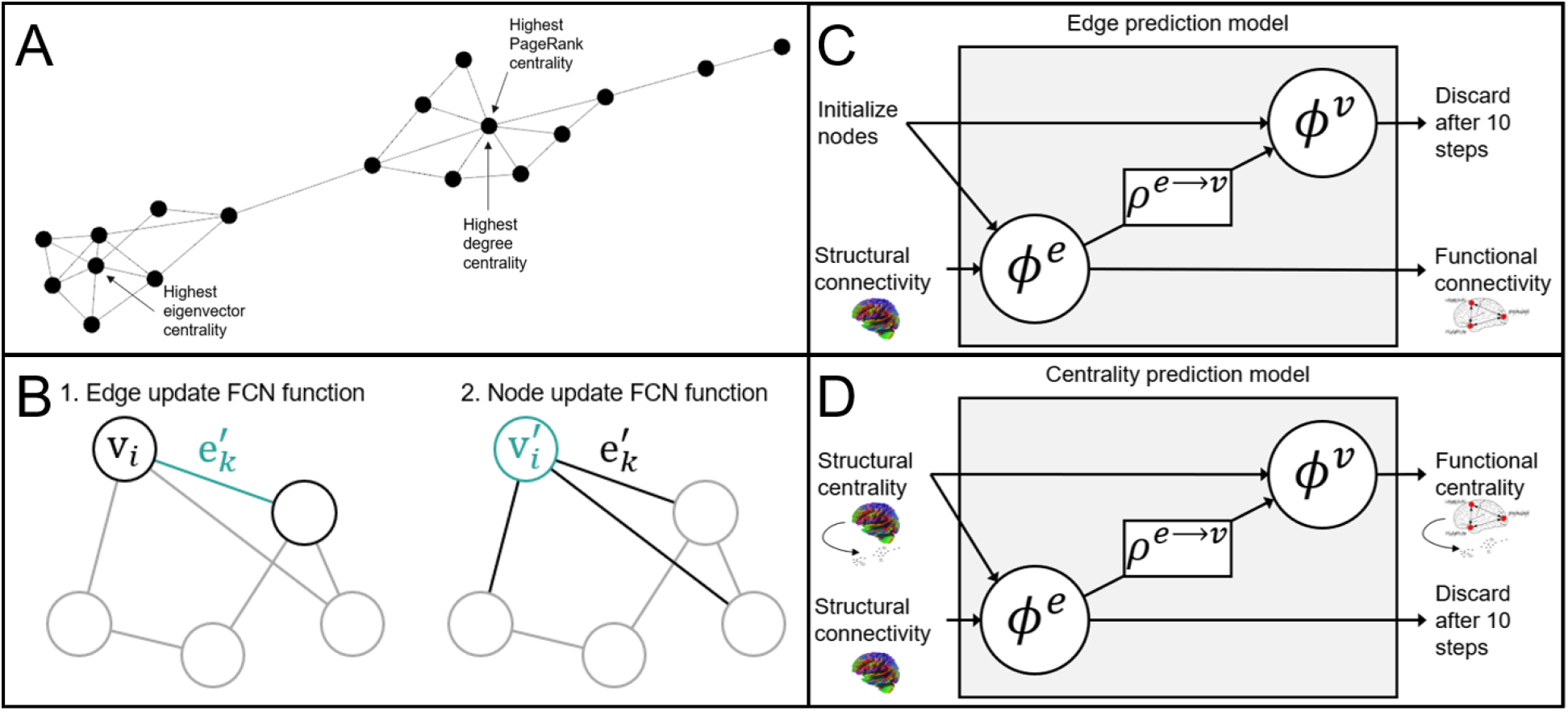
Graph neural network deep learning architecture. **(A)** An example graph illustrating degree centrality, eigenvector centrality, and PageRank centrality. **(B)** Depiction of Graph Nets update functions, where 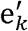 refers to the updated edge value, and where v_*i*_ and 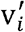 refer to the node being updated and then its updated value. Adapted from Battaglia et al. (2018). **(C and D)** Depiction of the steps in the edge prediction model **(C)**, and centrality prediction model **(D)**, where *ϕ*^*e*^ represents the FCN update function for edges taking an edge and 2 connected nodes as input, *ϕ*^*v*^ represents the FCN update function for nodes taking a node and the aggregated value of connected edges as input, and *ρ*^*e*→*ν*^ represents the aggregation of edge values. Adapted from Battaglia et al. (2018).

The problem of developing a deep learning model approach to using brain connectivity for prediction has been a focus of recent research. In a graph neural network model for deep learning, the structure of the connectivity data as a network (graph) is maintained, making this model ideal for prediction problems related to connectivity data. One implementation of this model architecture (not using additional global values in this case), Graph Nets (Battaglia et al., 2018), trains a small fully connected network (FCN) edge update function to update the value of each edge. This edge update function takes as input the current value of the edge as well as the values of each connected node. Another trainable FCN node update function updates the value of each edge. This node update function takes as input the current value of the node as well as the aggregated value of all connected edges (see Figure 1B). These two FCN functions can produce very good prediction results even at a very constrained scale, given only two layers with 16 nodes each (many fewer parameters to train than previous deep learning models predicting functional connectivity from structural connectivity; e.g., 4 layers with 350 nodes each in Rosenthal et al., 2018; 8 layers with 1024 nodes each in Sarwar et al., 2021), owing to the ability of the model to preserve the network structure of the data. Any number of updates can be performed, each using the updated values from the last step, before calculating the loss function and training the update functions (message passing). Graph neural network approaches to deep learning have been consistently outperforming other models of deep learning with brain connectivity data, as evidenced by higher prediction accuracy. Some applications of a graph neural network model to brain connectivity data include demonstrations of sex prediction from functional connectivity (88% accuracy; Arslan et al., 2018), a similarity metric learning model for predicting the similarity between two functional connectivity networks (63% accuracy; Ktena et al., 2017), prediction of Alzheimer’s disease from functional connectivity that outperformed previous methods (81% accuracy; Bi et al., 2020; Parisot et al., 2018), and prediction of Autism Spectrum Disorder outperforming previous methods (61% to 71%; Arya et al., 2020; Parisot et al., 2018; Wang et al., 2021; Zhang and Wang, 2020). To understand why graph neural network approaches have consistently outperformed other deep learning methods in neuroscience, it is important to understand that unlike FCN methods used previously, the graph neural network approach maintains the network structure of structural and functional connectivity. Conversely, the drawback of a FCN method is that all edge values are reduced to a single dimensional vector, which discards the vital network level patterns of connectivity. By maintaining the network structure, the graph neural network approach is able to produce superior performance using a much more constrained architecture with many fewer parameters (an edge and a node update function with only 2 layers of 16 nodes each, compared to FCN methods for example with 8 layers of 1024 nodes each). Additionally, because the graph neural network update functions apply the same function for every edge and node, this model applies a graph theory function that may represent a plausible explicit structure-function relationship to be explored with future research.

Considering the ultimate goal of relating DTI structural connectivity to resting state functional connectivity, a graph neural network deep learning approach has not yet been established, and based on past successes predicting other measures using brain connectivity this type of model is a promising candidate. This research uses a Graph Nets (Battaglia et al., 2018) deep learning model to determine to what extent structural connectivity can predict functional connectivity, as well as functional connectivity derived centrality measures.

## Methods

### Dataset

High quality DTI and resting state fMRI data for 998 subjects were obtained from the Human Connectome Project (HCP) database (Van Essen et al., 2013; please see this paper for ethics statements). The HCP preprocessed rsfMRI data was used, which has been FSL FIX (Salimi-Khorshidi et al., 2014) preprocessed, along with the DTI preprocessed data (see HCP preprocessing pipelines for more information on preprocessing steps, Glasser et al., 2013). The mean activation at each timepoint was calculated for each region of the Desikan-Killiany 66 region atlas (DK; Desikan et al., 2006; removed the corpus collosum region as in recent updates; e.g., Destrieux et al., 2010; Klein and Tourville, 2012) and of the Automated Anatomical Labelling 90 region atlas (AAL; Tzourio-Mazoyer et al., 2002). The DK atlas was used in the previous best prediction attempt (Sarwar et al., 2021), and multiple atlases were examined as the structure-function relationship is known to be atlas dependent (Messé, 2020). Each of the 4 rsfMRI sessions was then z-score standardized for each region independently from each other session. Each region was then bandpass filtered for each session separately to keep frequencies between 0.01 Hz and 0.1 Hz (see Hallquist et al., 2013).

### Connectivity Measures

The Pearson correlation coefficient was then calculated for each pair of regions using the 4800 total time points (all 4 sessions concatenated), as a measure of functional connectivity. DSI Studio’s (http://dsi-studio.labsolver.org) deterministic tracking algorithm that uses quantitative anisotropy (Yeh et al., 2013) as the termination index was used to produce structural connectivity matrices of streamline count. For reconstruction in DSI Studio the Generalized Q-sampling (Yeh et al., 2010) method was used, and tracking was performed using a fiber count of 1 million fibers, maximum angular deviation of 75 degrees, and a minimum and maximum fiber length of 20 mm and 500 mm respectively. A whole brain seed was used to calculate the structural connectivity matrix as the count of streamlines between each combination of the atlas regions.

Graph theory centrality measures of degree centrality, eigenvector centrality, and PageRank centrality were calculated using the NetworkX python library (Hagberg et al., 2008; using functions *degree, eigenvector_centrality*, and *pagerank*). These measures were calculated using a thresholded functional connectivity matrix, with the lower threshold set to the critical correlation coefficient for R with p = .0001 (see Zuo et al., 2012). Likewise, the DTI structural connectivity matrix was used to calculate degree, eigenvector, and PageRank centrality for each atlas region. The structural and functional centrality measures were then z-score standardized and rescaled to have values between -1 and 1.

### Model Architecture

The Graph Nets (Battaglia et al., 2018) python library, which relies on Tensorflow (Abadi et al., 2015), was used with a message passing design, using 10 message passing steps (as in the default model defined by Battaglia et al., 2018; accessible at https://github.com/albertotono/graph_nets; the code used to apply this library is available upon reasonable request). Following Battaglia et al., (2018) two latent layers were used with 16 nodes in each layer for the update functions. Seventy percent of the data was used as a training set, with 10 percent used for validation during training and 20 percent saved for testing (as done previously; e.g., Wang et al., 2020). The Adam (Kingma and Ba, 2017) optimizer was used with default values (learning rate = .001, β_1_ = .9, β_2_ = .999, and ε = 1 * 10^−7^). Training was performed using batch gradient descent, with a batch size of 32 used for training the models. The loss function used was the mean squared error (MSE) of the functional measures compared to the predicted values. One epoch was completed when enough batches were completed to randomly sample as much unique test data as possible (21 batches for batch size of 32). For the edge prediction models, structural connectivity values were given as the input edges, all input nodes were initialized with a value of 1.0, and functional connectivity measures were compared to the output edges to calculate loss (see Figure 1C). For the centrality prediction models, structural connectivity values were given as the input edges, structural centrality measures were given as the input nodes, and functional centrality measures were compared to the output nodes to calculate loss (see Figure 1D). In order to encourage the model to find an effective solution in as few steps as possible, the loss was calculated as the mean MSE for all 10 message passing steps. A Nvidia GTX 3080 graphical processing unit (GPU) was used to train the models and required approximately 11 hours to run 3000 epochs for the centrality prediction models and approximately 83 hours to run 5000 epochs for the edge prediction model.

## Results

### Edges

The edge prediction model was trained for 5000 epochs, and there was no evidence for overfitting of the model to the training set, as the validation set decreased its loss along with the training set (Figures 2A and 3A). Figures 2B and 3B depict the empirical versus the mean predicted values of functional connectivity, accounting for 89.4% (DK) and 81.3% (AAL) of the variance in the 201 subject test group. Furthermore, functional connectivity was predicted at the individual-level accounting for 55.7% (DK) and 47.8% (AAL) of the variance (see Supplementary Figures 1 and 2). The structural connectivity of each individual atlas region was iteratively lesioned, and this lesioned network was given to the previously trained model as input. The MSE lesioned loss was then calculated, whereby an increase in loss following a removal of that region indicates the importance of that region to model performance (see Figures 2C and 3C). For the DK atlas the top 10 regions that impacted the model performance the most included bilateral superior frontal cortex, bilateral precentral gyrus, superior parietal lobe, insula, and lingual gyrus. For the AAL atlas the top 10 regions that impacted the model performance the most included the bilateral middle occipital gyrus, RH angular gyrus, LH precentral gyrus, RH superior frontal lobe, RH middle cingulum, RH olfactory cortex, LH insula, LH middle temporal pole, and RH lingual gyrus. In order to gain a better understanding of what features of the regions make them important for model performance, we tested a hypothesis that higher structural degree centrality (more centrally important to the network as a whole because of the high number of connections) may be related to higher impact on model performance. This was the case, as it was observed that the centrality of a region was positively related to lesion loss (Figures 2D and 3D; DK: *R*(64) = .774, *p* < .001; AAL: *R*(88) = .322, *p* = .002), indicating that high centrality regions were more important for model performance (see Figure 2D).

**Figure 2.**
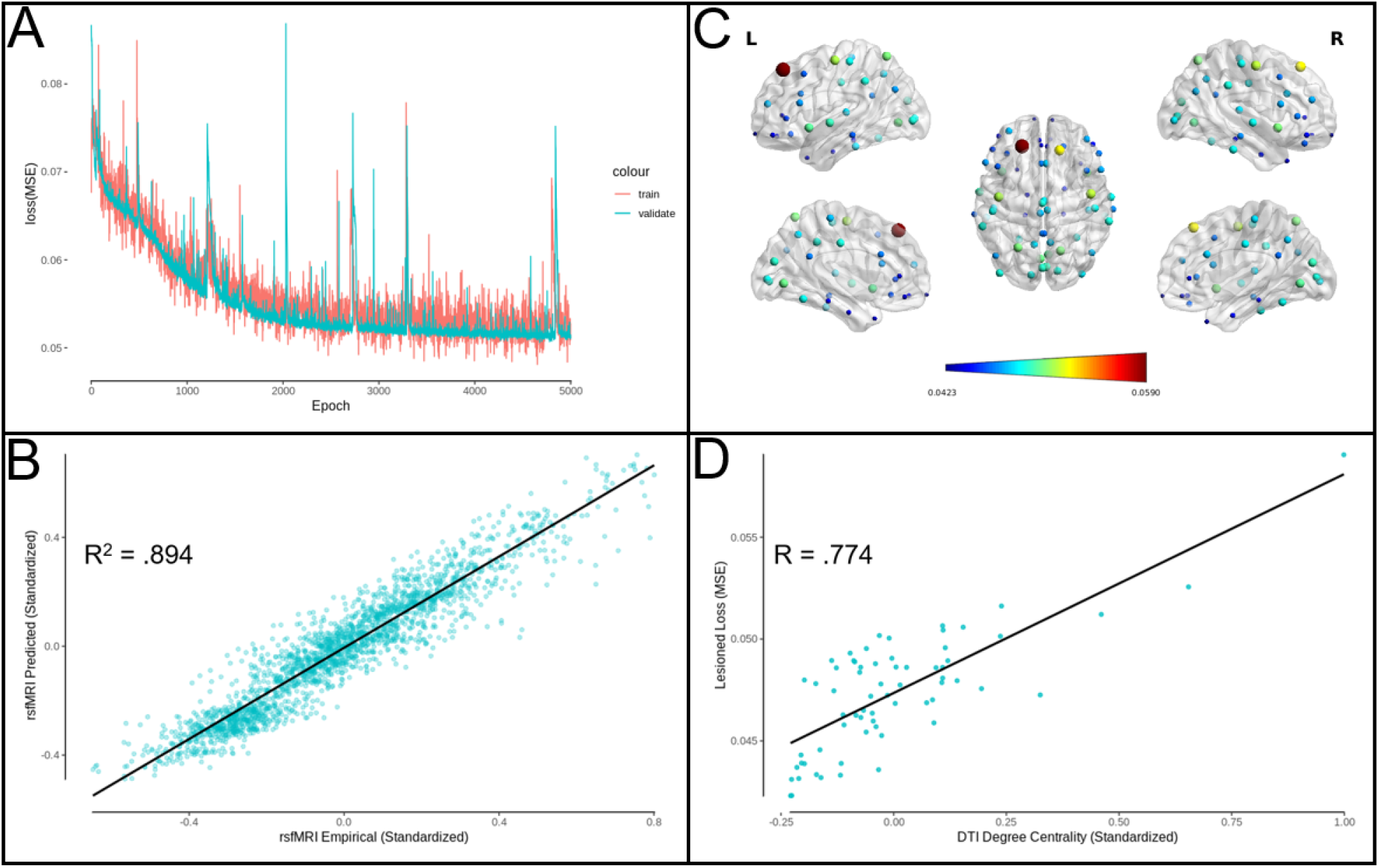
DK atlas edge prediction model performance. **(A)** Edge prediction model MSE loss for training and validation sets as a function of the number of training epochs. **(B)** Predicted rsfMRI functional connectivity as a function of empirical rsfMRI functional connectivity (R^2^ = .894). **(C)** Functional connectivity loss (MSE) related to lesioning structural connectivity to each atlas region, where dark blue indicates a lesser effect on the model performance and dark red indicates a greater effect. Larger sphere size also indicates a greater effect of lesion on model performance. Figure produced using BrainNet Viewer (Xia et al., 2013). **(D)** Lesioned functional connectivity loss as a function of structural connectivity degree centrality (*R*(64) = .774, *p* < .001).

**Figure 3.**
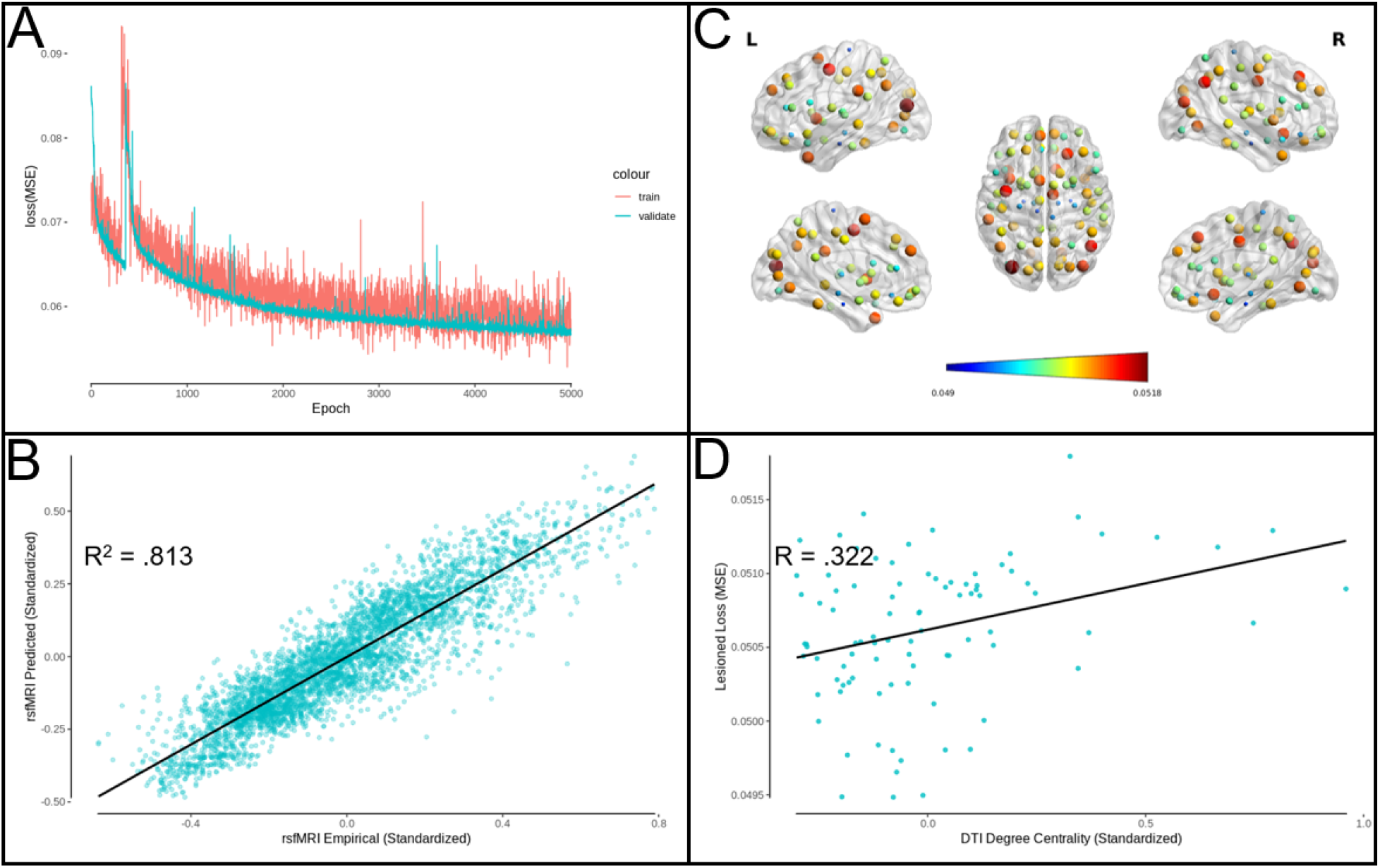
AAL atlas edge prediction model performance. **(A)** Edge prediction model MSE loss for training and validation sets as a function of the number of training epochs. **(B)** Predicted rsfMRI functional connectivity as a function of empirical rsfMRI functional connectivity (R^2^ = .813). **(C)** Functional connectivity loss (MSE) related to lesioning structural connectivity to each atlas region, where dark blue indicates a lesser effect on the model performance and dark red indicates a greater effect. Larger sphere size also indicates a greater effect of lesion on model performance. Figure produced using BrainNet Viewer (Xia et al., 2013). **(D)** Lesioned functional connectivity loss as a function of structural connectivity degree centrality (*R*(88) = .322, *p* = .002).

Using the predicted functional connectivity, measures of centrality for degree, eigenvector, and PageRank were calculated by unstandardizing first before following the same thresholding and centrality calculation as described in the Methods. The mean predicted values for degree centrality accounted for 88.2% (DK) and 80.8% (AAL) of the variance in the empirical data (see Figures 4A and 5A), for eigenvector centrality accounted for 89.9% (DK) and 84.0% (AAL) of the variance in the empirical data (see Figures 4B and 5B), and for PageRank centrality accounted for 88.9% (DK) and 80.0% (AAL) of the variance in the empirical data (see Figures 4C and 5C). The individual-level predicted values for degree centrality accounted for 81.0% (DK) and 73.1% (AAL) of the variance in the empirical data (see Supplementary Figures 3 and 4), for eigenvector centrality accounted for 55.3% (DK) and 53.0% (AAL) of the variance in the empirical data (see Supplementary Figures 5 and 6), and for PageRank centrality accounted for 55.0% (DK) and 50.8% (AAL) of the variance in the empirical data (see Supplementary Figures 7 and 8). These results demonstrate that the model has accounted for a large amount of variance in connectivity as well as centrality, but in order to determine if even better performance could be produced, the model designed to directly predict centrality was utilized.

**Figure 4.**
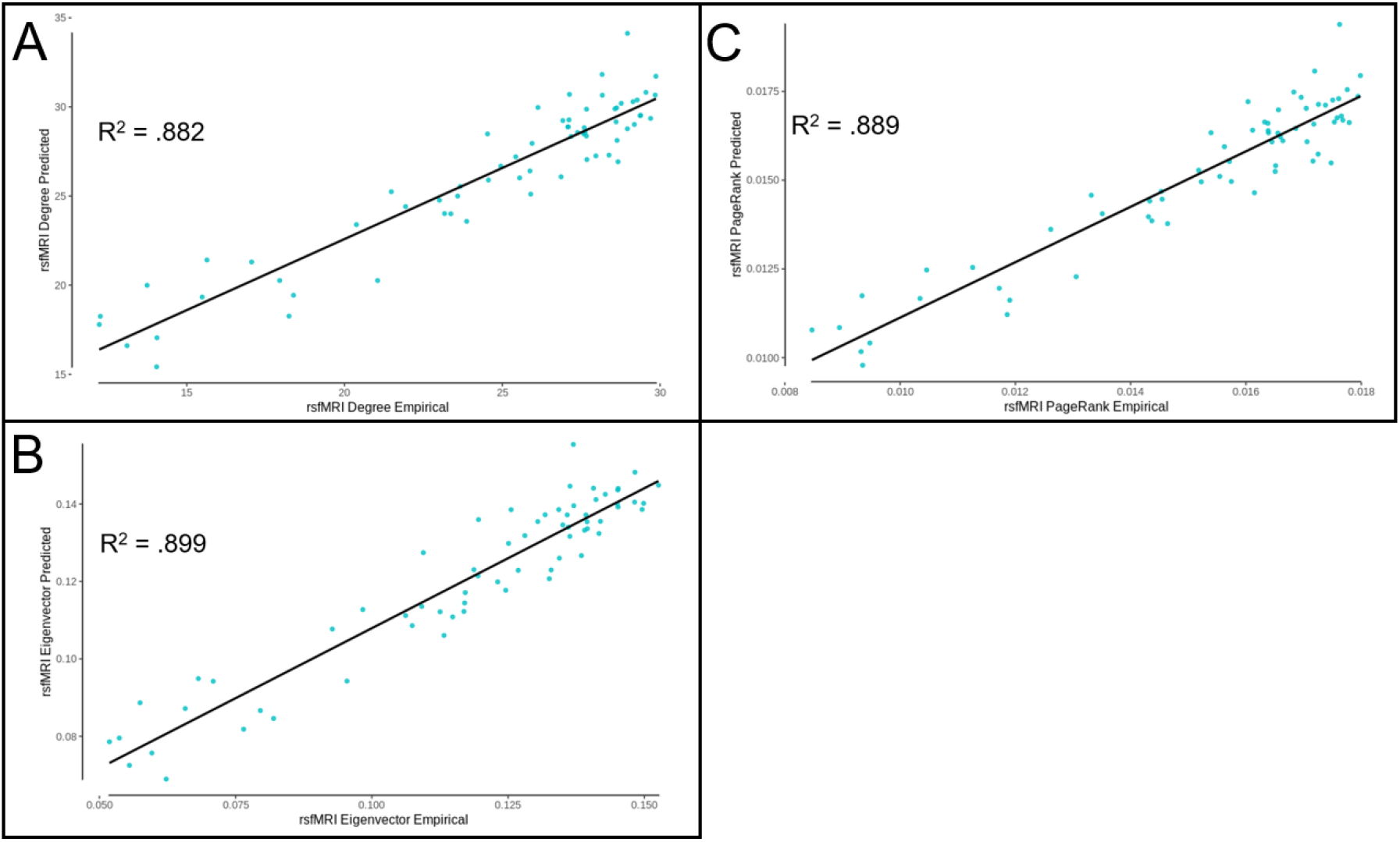
DK atlas edge prediction model derived centrality measure performance. **(A)** Degree centrality measures calculated from the predicted functional connectivity values as a function of empirical degree centrality (R^2^ = .882). **(B)** Eigenvector centrality measures calculated from the predicted functional connectivity values as a function of empirical eigenvector centrality (R^2^ = .899). **(C)** PageRank centrality measures calculated from the predicted functional connectivity values as a function of empirical PageRank centrality (R^2^ = .889).

**Figure 5.**
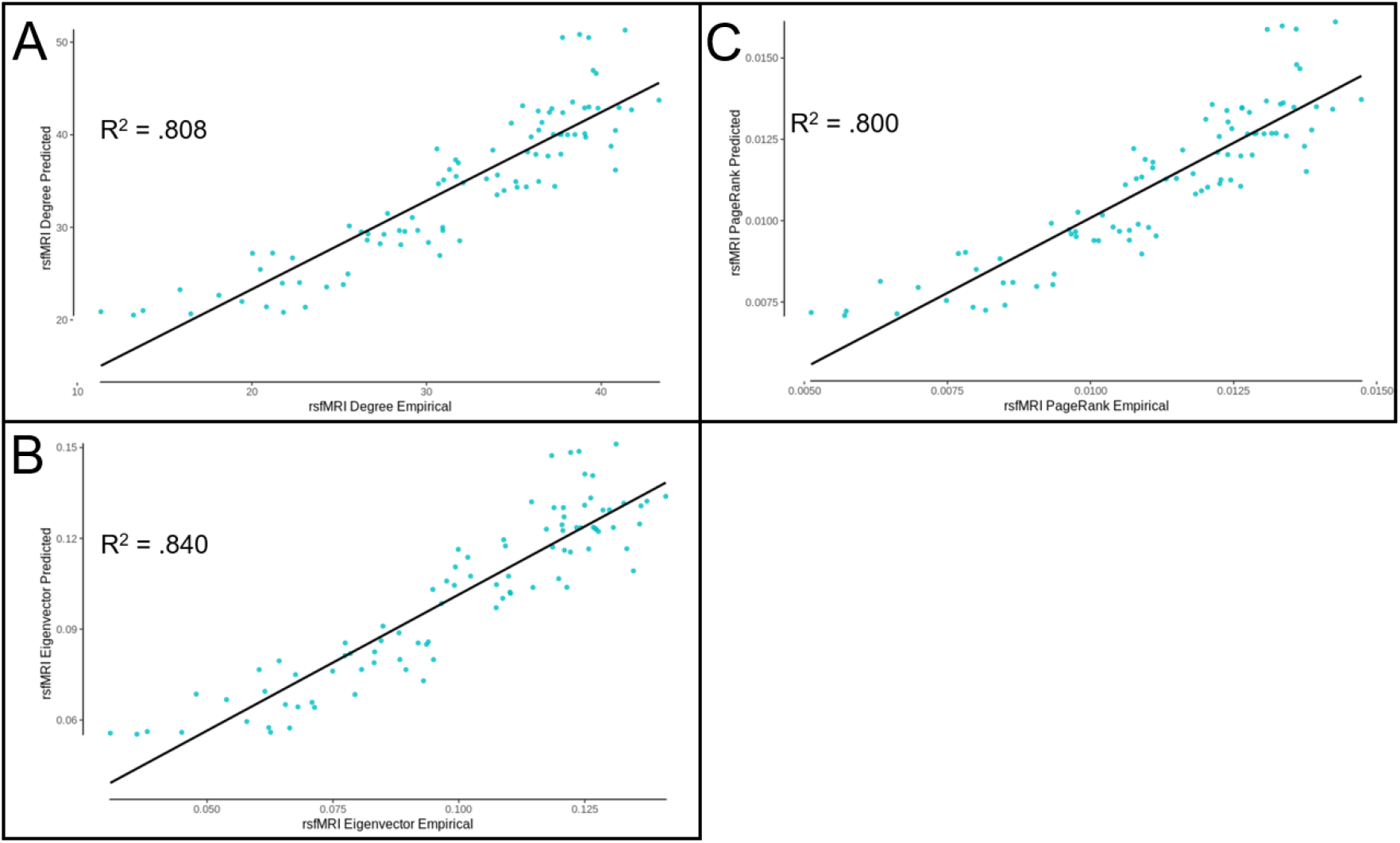
AAL atlas edge prediction model derived centrality measure performance. **(A)** Degree centrality measures calculated from the predicted functional connectivity values as a function of empirical degree centrality (R^2^ = .808). **(B)** Eigenvector centrality measures calculated from the predicted functional connectivity values as a function of empirical eigenvector centrality (R^2^ = .840). **(C)** PageRank centrality measures calculated from the predicted functional connectivity values as a function of empirical PageRank centrality (R^2^ = .800).

### Degree Centrality

The degree prediction model was trained for 3000 epochs, and there was no evidence for overfitting of the model to the training set, as the validation set decreased its loss along with the training set (Figures 6A and 7A). Figures 6B and 7B depict the empirical versus the predicted mean values of functional degree centrality, accounting for 99.3% (DK) and 99.0% (AAL) of the variance in the 201 subject test group. The individual-level functional degree centrality was predicted accounting for 63.7% (DK) and 64.7% (AAL) of the variance (Supplementary Figures 9 and 10). This alternative model performed better than the previous model predicting the mean centrality, which accounted for 88.2% (DK) and 80.8% (AAL) of the variance, but performed worse than the previous model predicting the individual-level centrality, which accounted for 81.0% (DK) and 73.1% (AAL) of the variance.

**Figure 6.**
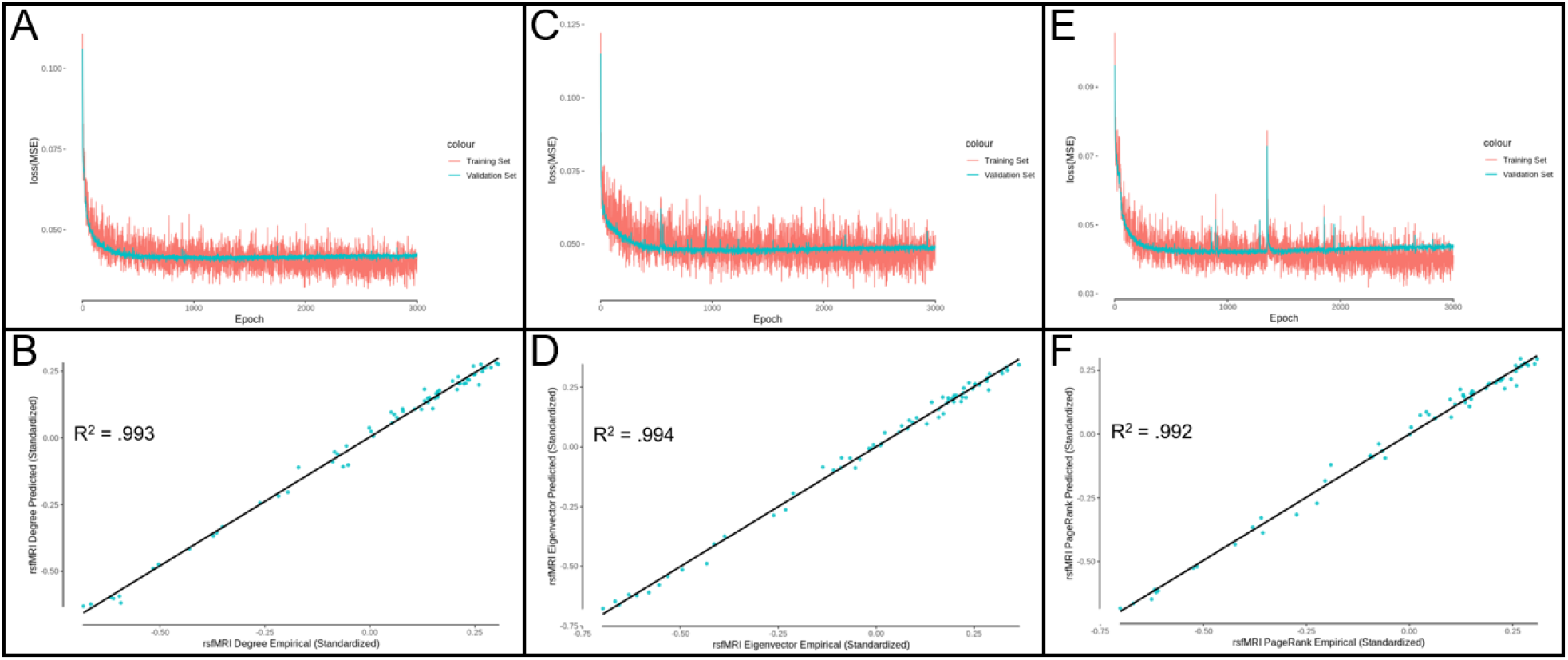
DK atlas centrality prediction model performance. **(A)** Degree centrality model MSE loss for training and validation sets as a function of the number of training epochs. **(B)** Predicted rsfMRI functional connectivity degree centrality as a function of empirical rsfMRI functional connectivity degree centrality (R^2^ = .993). **(C)** Eigenvector centrality model MSE loss for training and validation sets as a function of the number of training epochs. **(D)** Predicted rsfMRI functional connectivity eigenvector centrality as a function of empirical rsfMRI functional connectivity eigenvector centrality (R^2^ = .994). **(E)** PageRank centrality model MSE loss for training and validation sets as a function of the number of training epochs. **(F)** Predicted rsfMRI functional connectivity PageRank centrality as a function of empirical rsfMRI functional connectivity PageRank centrality (R^2^ = .992).

**Figure 7.**
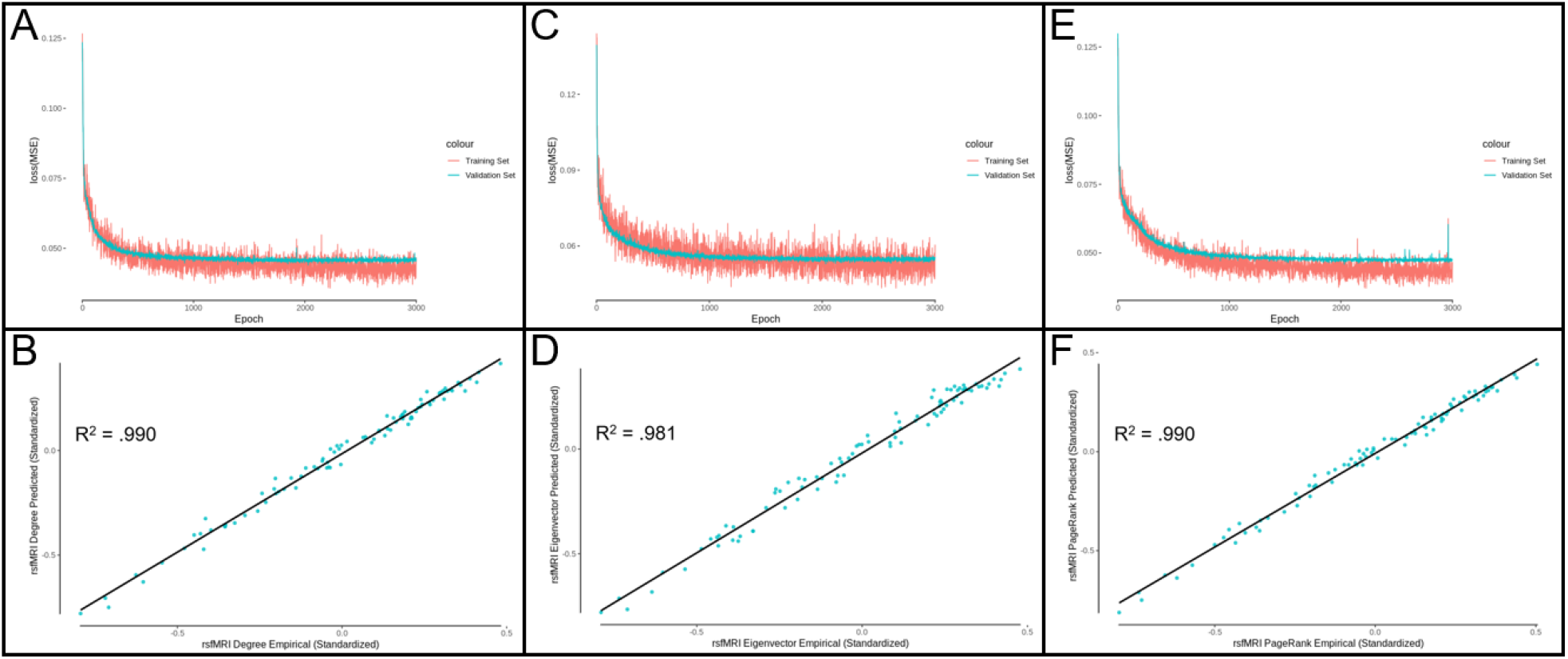
AAL atlas centrality prediction model performance. **(A)** Degree centrality model MSE loss for training and validation sets as a function of the number of training epochs. **(B)** Predicted rsfMRI functional connectivity degree centrality as a function of empirical rsfMRI functional connectivity degree centrality (R^2^ = .990). **(C)** Eigenvector centrality model MSE loss for training and validation sets as a function of the number of training epochs. **(D)** Predicted rsfMRI functional connectivity eigenvector centrality as a function of empirical rsfMRI functional connectivity eigenvector centrality (R^2^ = .981). **(E)** PageRank centrality model MSE loss for training and validation sets as a function of the number of training epochs. **(F)** Predicted rsfMRI functional connectivity PageRank centrality as a function of empirical rsfMRI functional connectivity PageRank centrality (R^2^ = .990).

### Eigenvector Centrality

The eigenvector prediction model was trained for 3000 epochs, and there was no evidence for overfitting of the model to the training set, as the validation set decreased its loss along with the training set (Figures 6C and 7C). Figures 6D and 7D depict the empirical versus the predicted mean values of functional eigenvector centrality, accounting for 99.4% (DK) and 98.1% (AAL) of the variance in the 201 subject test group. The individual-level functional eigenvector centrality was predicted accounting for 62.3% (DK) and 61.2% (AAL) of the variance (Supplementary Figures 11 and 12). This alternative model performed better than the previous model, which accounted for 89.9% (DK) and 84.0% (AAL) of the mean variance and 55.3% (DK) and 53.0% (AAL) of the individual-level variance.

### PageRank Centrality

The PageRank prediction model was trained for 3000 epochs, and there was no evidence for overfitting of the model to the training set, as the validation set decreased its loss along with the training set (Figures 6E and 7E). Figures 6F and 7F depict the empirical versus the predicted mean values of functional PageRank centrality, accounting for 99.2% (DK) and 99.0% (AAL) of the variance in the 201 subject test group. The individual-level functional PageRank centrality was predicted accounting for 64.0% (DK) and 64.9% (AAL) of the variance (Supplementary Figures 13 and 14). This alternative model performed better than the previous model, which accounted for 88.9% (DK) and 80.0% (AAL) of the mean variance and 55.0% (DK) and 50.8% (AAL) of the individual-level variance.

## Discussion

Resting state fMRI has been used to calculate functional connectivity networks in brain research for many years, and graph theory measures such as centrality have been calculated for these functional networks, yet a clear relationship between the physical structural connectivity derived values and what is assumed to be the ‘effective’ functional connectivity has been elusive. The most successful previous effort to predict mean functional connectivity from mean structural connectivity at the whole-brain level was able to account for 81% of the variance in functional connectivity, and individual-level functional connectivity was predicted from individual-level structural connectivity accounting for 30% of the variance. The importance of continuing efforts to improve model prediction performance has been recently highlighted (Suárez et al., 2020). By using the Graph Nets (Battaglia et al., 2018) deep learning model architecture, which is well suited to modeling network datasets, mean functional connectivity was predicted from individual-level structural connectivity accounting for 89% of the variance (surpassing the previous attempt, without relying on mean structural connectivity as input), and individual-level functional connectivity was predicted from individual-level structural connectivity accounting for 48% of the variance (far surpassing the previous attempt). In addition, mean functional centrality was predicted from individual-level structural connectivity and centrality data accounting for up to 99% of the variance, and up to 81.0% of individual-level functional centrality variance was accounted for from individual-level structural connectivity, demonstrating that these functional centrality measures can be robustly derived from the underlying structural connectivity and structural centrality measures. These results demonstrate that it is possible to account for nearly all of the mean-level variance in functional centrality with structural connectivity and centrality measures, suggesting that by calculating graph theory measures, information from the whole network is integrated, bridging the gap that is much more difficult to cross between structure and function at the edge level, and should encourage further research to explicitly define the nature of this relationship using graph theory and simulation modelling. Note that there is still variance left to be accounted for in predicting centrality at the individual level, which is not surprising considering that local structural connectivity at the microscale very likely contributes to functional centrality in a way that cannot be captured using macroscale methods. These results set an important benchmark in what should be a continuing effort in the network neuroscience community to determine the extent to which functional connectivity and graph theory measures such as centrality can be derived from structural connectivity, and therefore to what extent we can infer functional connectivity from structural connectivity.

The regions of the brain that are particularly important for the performance of the edge prediction model were highlighted by iteratively lesioning each region. In order to test whether the centrality may affect how important a region is to model performance, degree centrality was compared to lesion loss. Indeed, higher centrality was associated with greater importance for model performance, indicating that part of what makes a region more important to the model is the extent to which the region is centrally important to the network, connected to many other regions. As with all deep learning approaches, the limitations of this model include the nonlinear, underlying function that acts on the structural connectivity to predict the functional connectivity. Although we have addressed this limitation in part by showing a brain map of the regional importance to model performance via iterative lesioning, future research is needed using explicit graph theory measures and simulation models to define the nature of the structure-function relationship.

### Future Directions

The graph neural network deep learning model architecture is one that is designed with networks in mind, and maintains the structure of the brain connectivity data, whereas other commonly used deep learning approaches do not. We have demonstrated that this is an effective deep learning model for use with structural and functional connectivity data, whether the edge-, node-, or participant-level measures are of interest. Future research predicting patient, demographic, or behavioural data should also make use of this method in order to improve on past deep learning attempts using the “global” participant-level output available for training in the graph neural network model. Prediction of task activation from structural connectivity has also been explored in recent research (e.g., Ekstrand et al., 2020; Osher et al., 2016; Wu et al., 2020), and this model could also lead to improvements in task activation prediction.

As this model is further developed to account for more variance in individual-level functional connectivity, this approach may also lead to predictive functional mapping for patients who are unable to follow fMRI instructions. Alternatively, there may be an upper limit on the amount of information about functional connectivity that is contained in structural connectivity measures. This upper limit may exist due to a number of factors. Neuromodulation selectively inhibits and excites neurons throughout the brain network, leading to dynamic patterns of functional connectivity, so although the underlying structure does not change during MRI imaging, the functional connectivity is in constant flux (Bell and Shine, 2016; Shine, 2019). The low spatial resolution of the atlas used also means that the precise structural connectivity patterns between neurons cannot be measured. In addition, functional boundaries can vary across individuals and between sessions (Gordon et al., 2017; Laumann et al., 2015; Mueller et al., 2013; Salehi et al., 2020; Suárez et al., 2020; Wang et al., 2015), so the use of atlases is a source of error for this reason as well. The temporal dynamics also differ from one region to the next, affecting the calculation of functional connectivity (Gollo et al., 2015; Keitel and Gross, 2016; Murray et al., 2014; Shafiei et al., 2019; Suárez et al., 2020). Depending on how unimodal or transmodal a region is, there may be a higher (unimodal) or lower (transmodal) degree of coupling between structure and function (Margulies et al., 2016; Preti and Van De Ville, 2019; Suárez et al., 2020). Based on our deep learning model results, there is clearly a large amount variance in functional connectivity that can be accounted for by structural connectivity and although the relationship discovered by deep learning is difficult to interpret directly, further graph theory and simulation modeling research can define this relationship explicitly, accounting for increased variance until deep learning-like levels of variance are accounted for.

## Conclusion

Our graph neural network deep learning model provides a new benchmark for prediction of rsfMRI functional connectivity (89% of mean variance; 56% of individual-level variance) and functional centrality (99% of mean variance; 81% of individual-level variance) from DTI structural connectivity, which far exceeds the individual-level performance of the non graph neural network models previously reported by others (81% of mean variance; 30% of individual-level variance). This research has not only brought us closer to finding the upper limit on prediction of connectivity and centrality of function from structural connectivity, it has opened new doors for understanding the structure-function network relationships in the human brain.

## Declarations

### Availability of data and material

All data is made publicly available by the Human Connectome Project (Van Essen et al., 2013).

### Code availability

The Graph Nets (Battaglia et al., 2018) python library is accessible at https://github.com/albertotono/graph_nets. The code used to apply this library is available upon reasonable request.

### Ethics approval, consent to participate, and consent for publication

Please see Van Essen et al. (2013) for a description of ethics and consent.

## Acknowledgements

This research was supported by the Natural Sciences and Engineering Research Council of Canada through Alexander Graham Bell Canada Graduate Scholarships to Josh Neudorf and Shaylyn Kress, and Discovery Grant 18968-2013-22 to the senior author Ron Borowsky. The authors affirm that there are no conflicts of interest to disclose.

Data were provided [in part] by the Human Connectome Project, WU-Minn Consortium (Principal Investigators: David Van Essen and Kamil Ugurbil; 1U54MH091657) funded by the 16 NIH Institutes and Centers that support the NIH Blueprint for Neuroscience Research; and by the McDonnell Center for Systems Neuroscience at Washington University.

## Supplementary Material

**Supplementary Figure 1.**
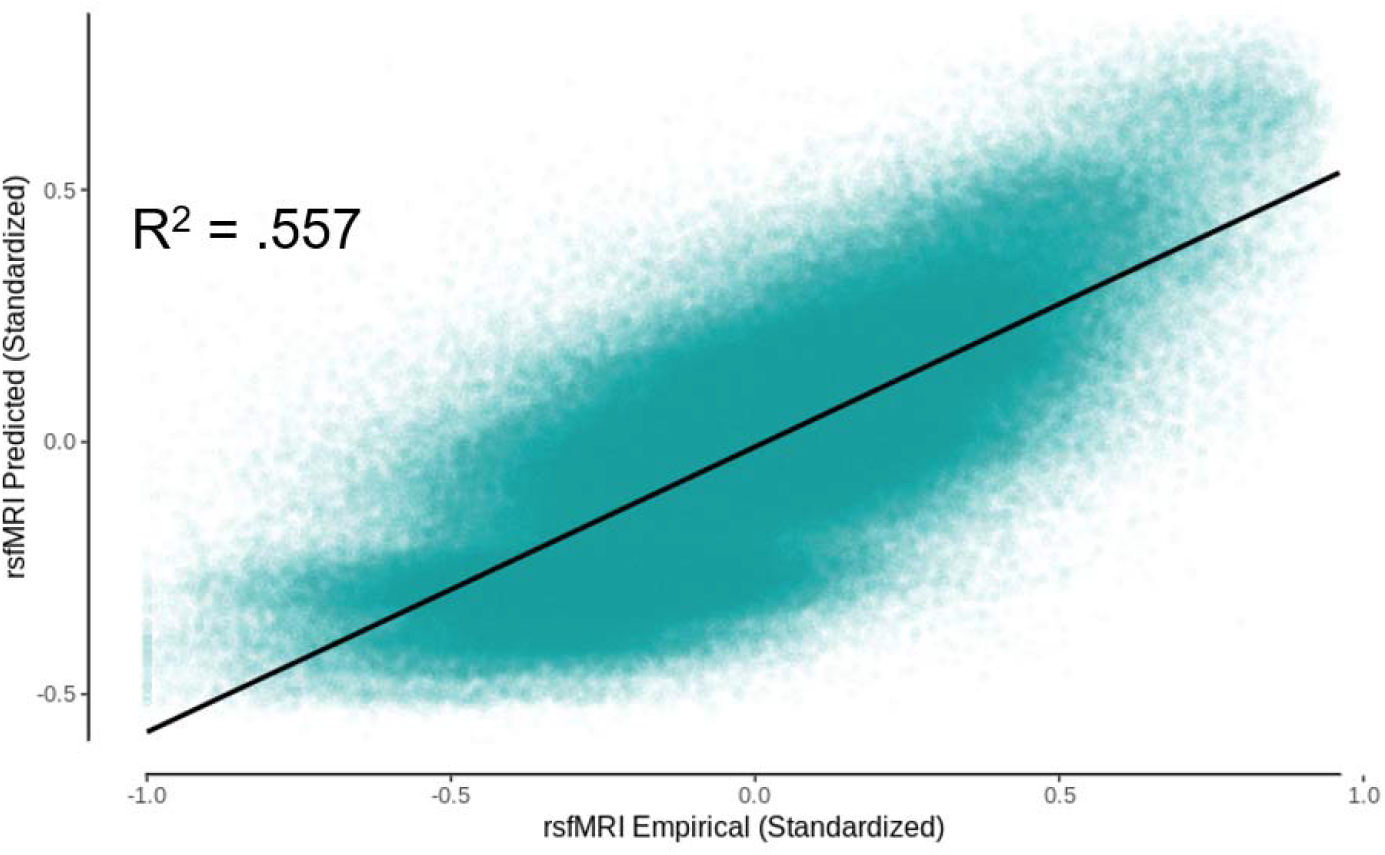
DK atlas non-aggregated predicted rsfMRI functional connectivity as a function of empirical rsfMRI functional connectivity (R^2^ = .557).

**Supplementary Figure 2.**
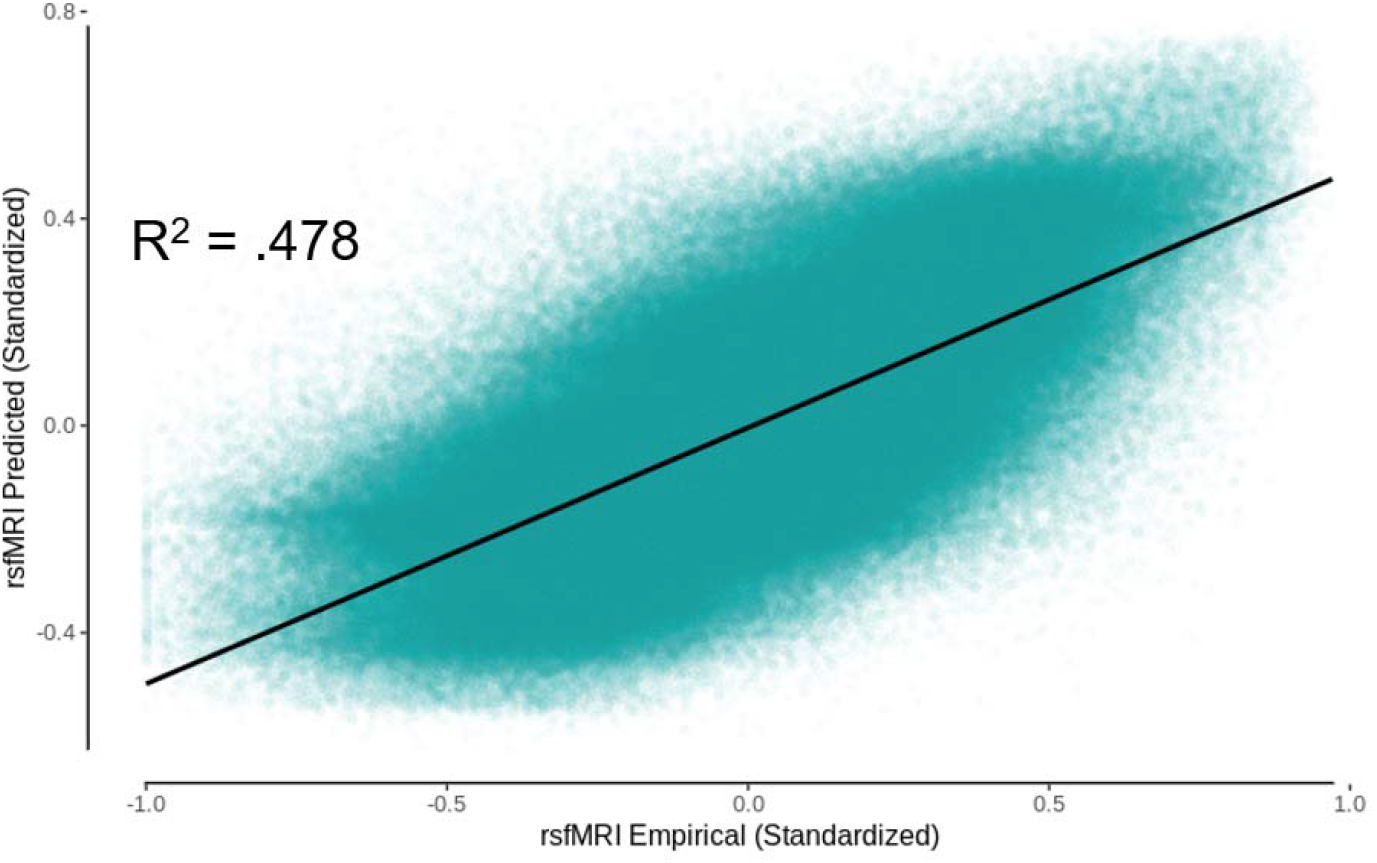
AAL atlas non-aggregated predicted rsfMRI functional connectivity as a function of empirical rsfMRI functional connectivity (R^2^ = .478).

**Supplementary Figure 3.**
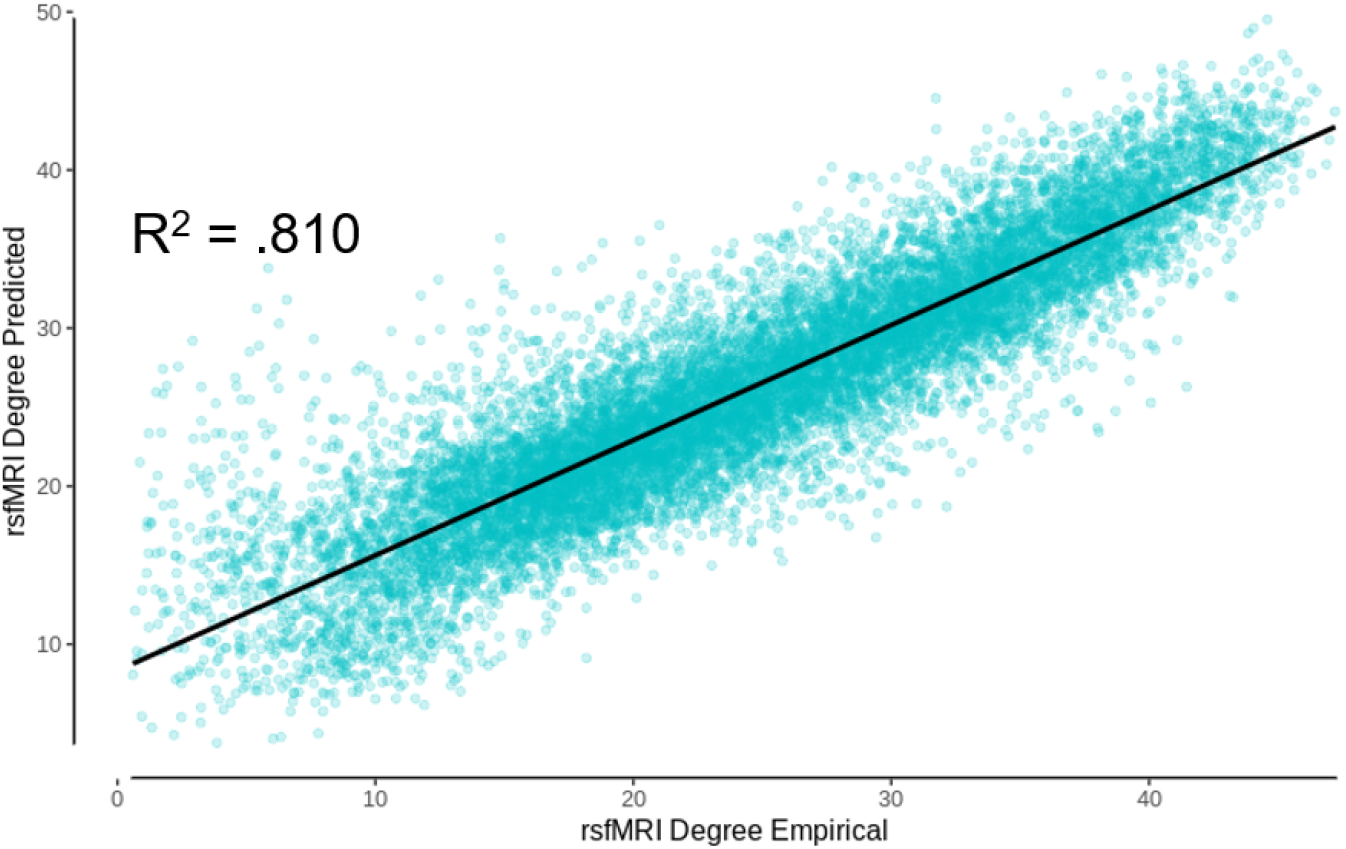
DK atlas non-aggregated degree centrality measures calculated from the predicted rsfMRI functional connectivity values as a function of empirical rsfMRI functional degree centrality (R^2^ = .810).

**Supplementary Figure 4.**
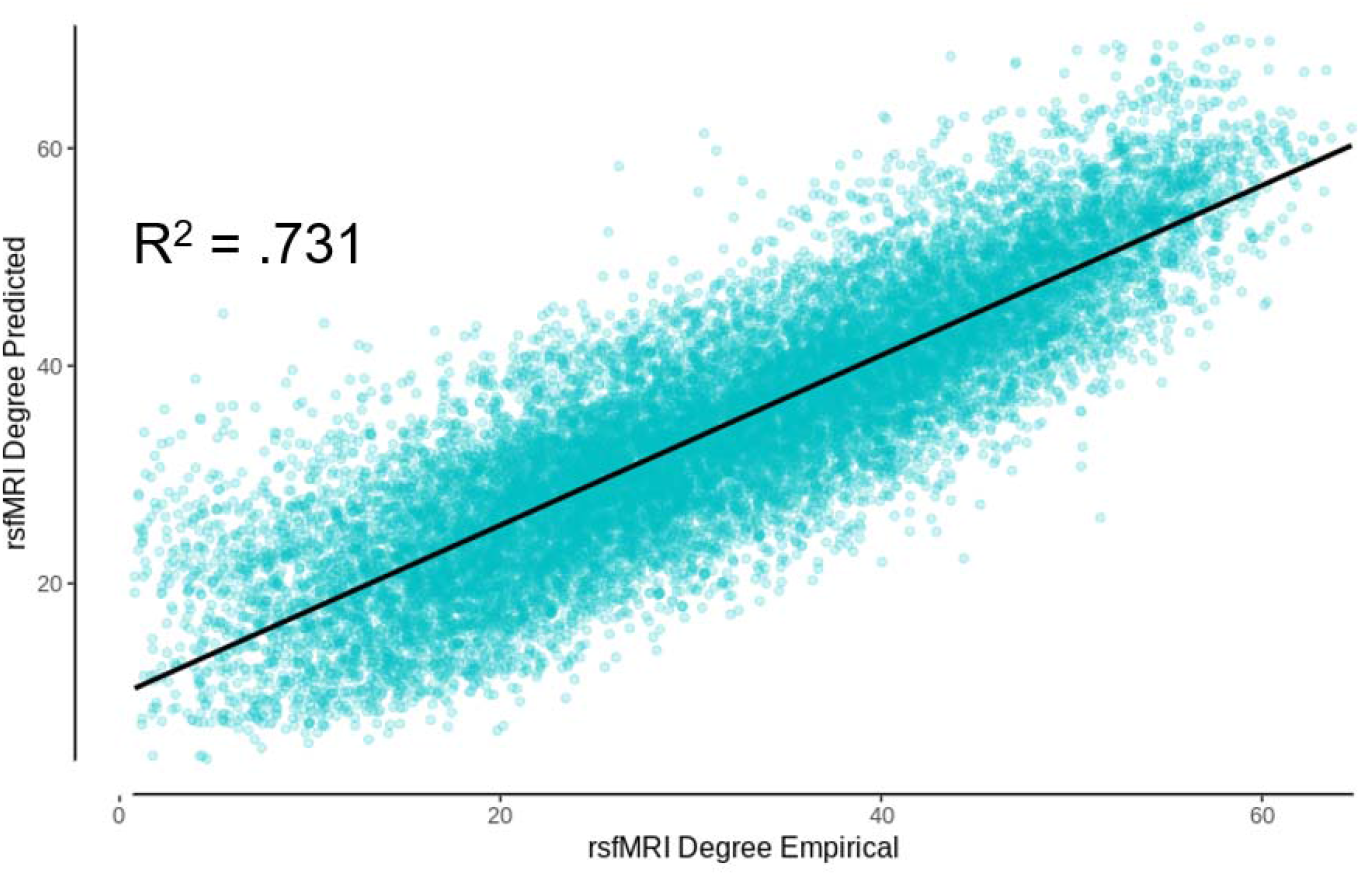
AAL atlas non-aggregated degree centrality measures calculated from the predicted rsfMRI functional connectivity values as a function of empirical rsfMRI functional degree centrality (R^2^ = .731).

**Supplementary Figure 5.**
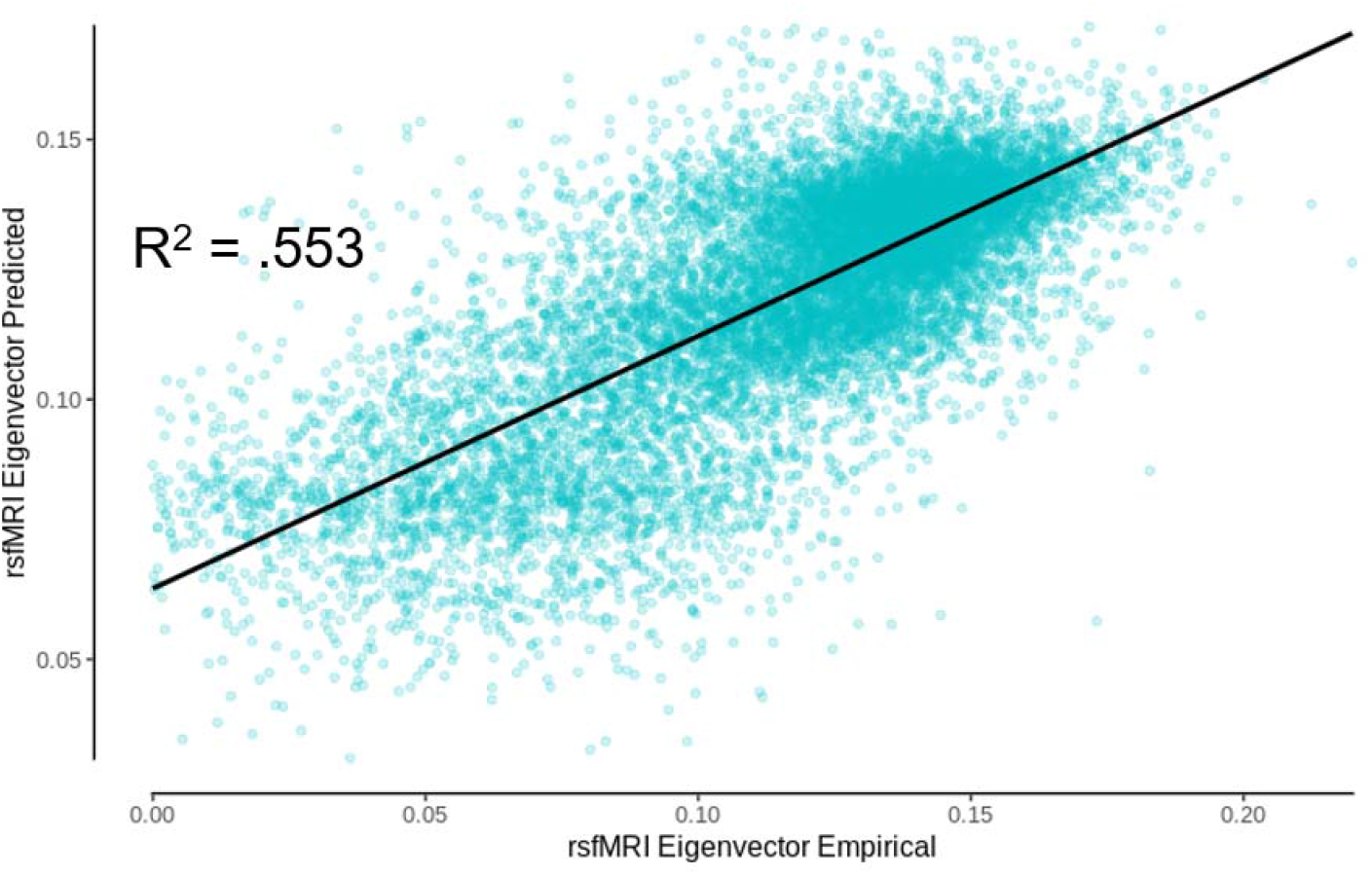
DK atlas non-aggregated rsfMRI functional eigenvector centrality measures calculated from the predicted functional connectivity values as a function of empirical rsfMRI functional eigenvector centrality (R^2^ = .553).

**Supplementary Figure 6.**
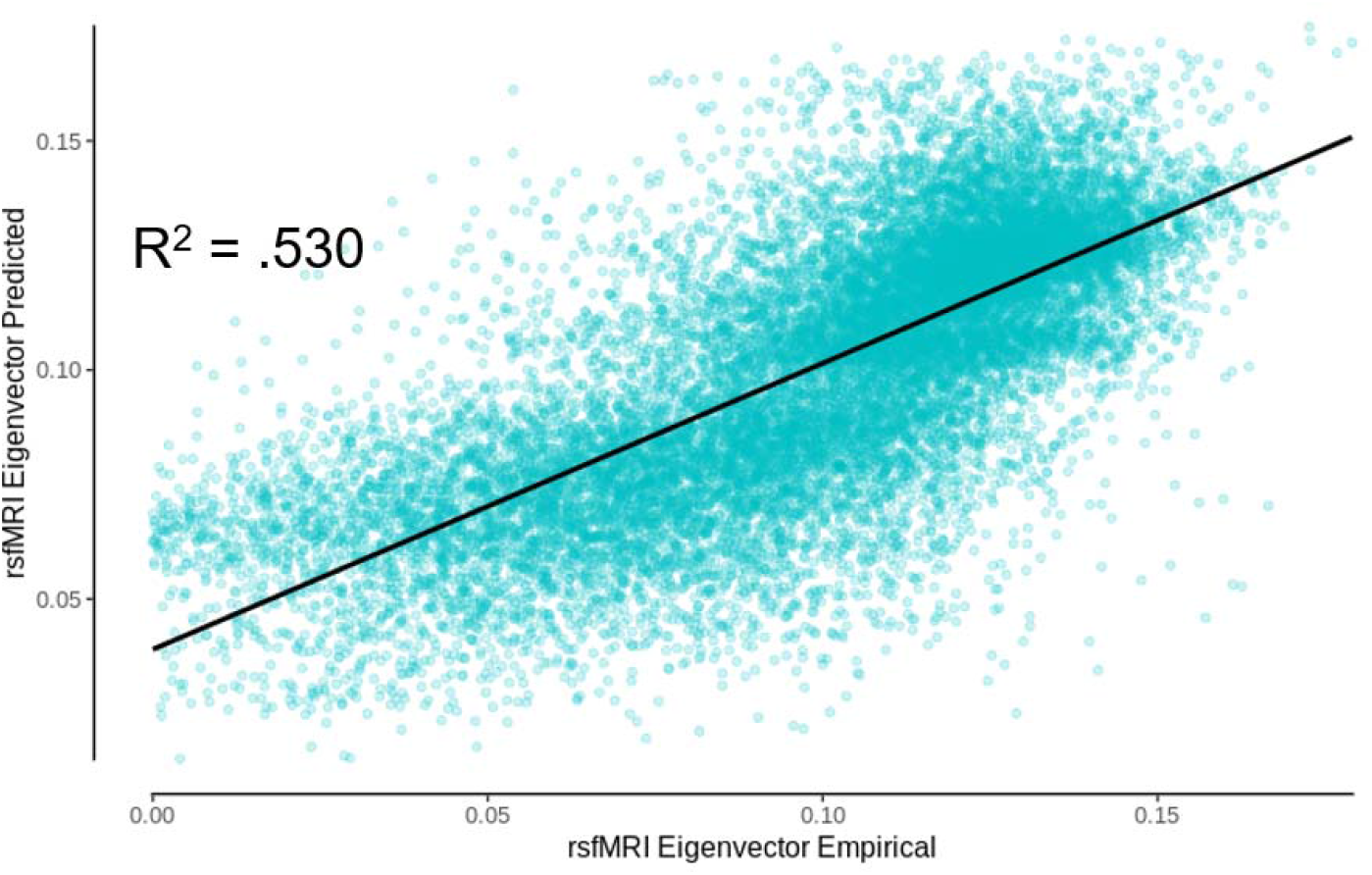
AAL atlas non-aggregated rsfMRI functional eigenvector centrality measures calculated from the predicted functional connectivity values as a function of empirical rsfMRI functional eigenvector centrality (R^2^ = .530).

**Supplementary Figure 7.**
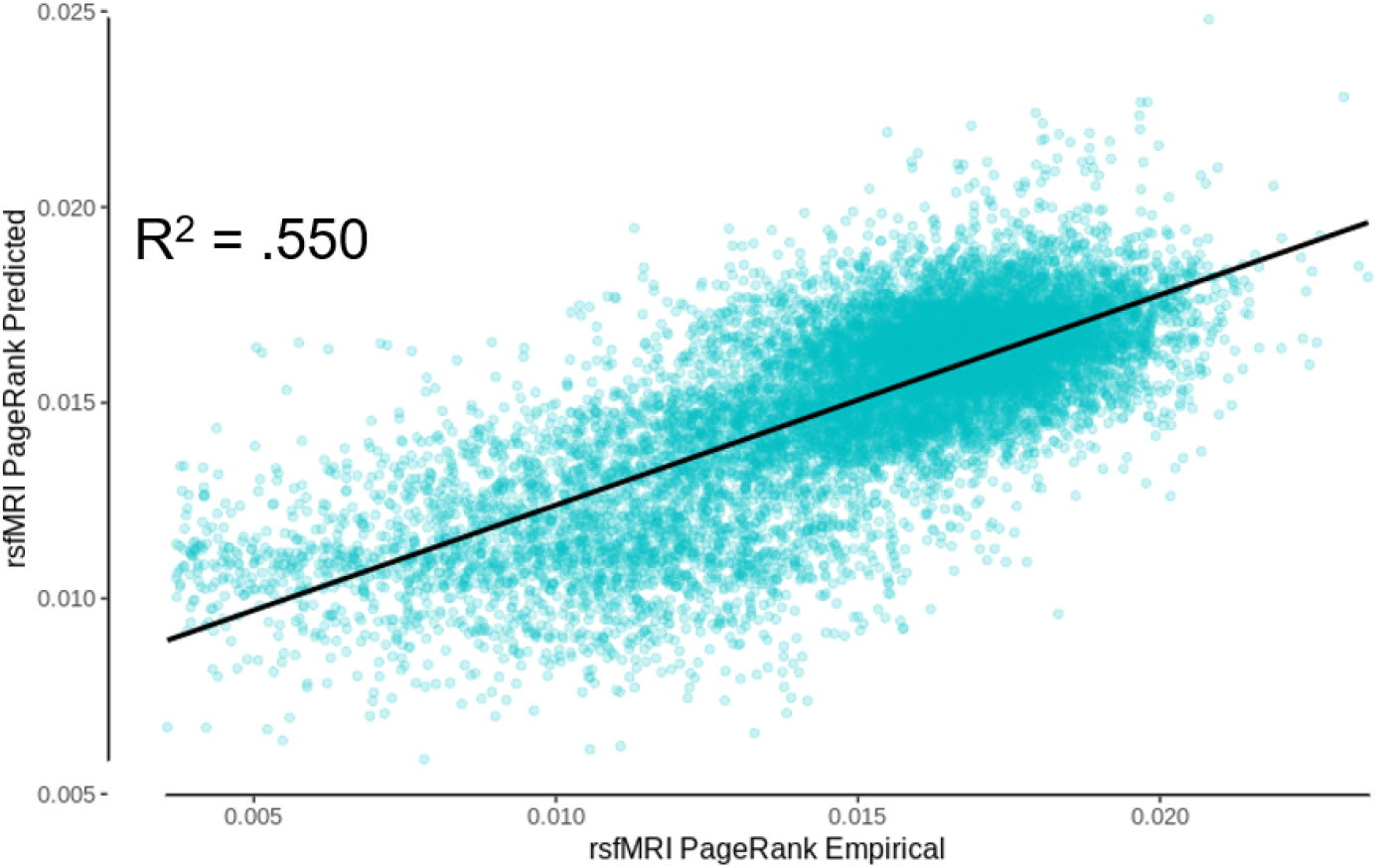
DK atlas non-aggregated rsfMRI functional PageRank centrality measures calculated from the predicted functional connectivity values as a function of empirical rsfMRI functional PageRank centrality (R^2^ = .550).

**Supplementary Figure 8.**
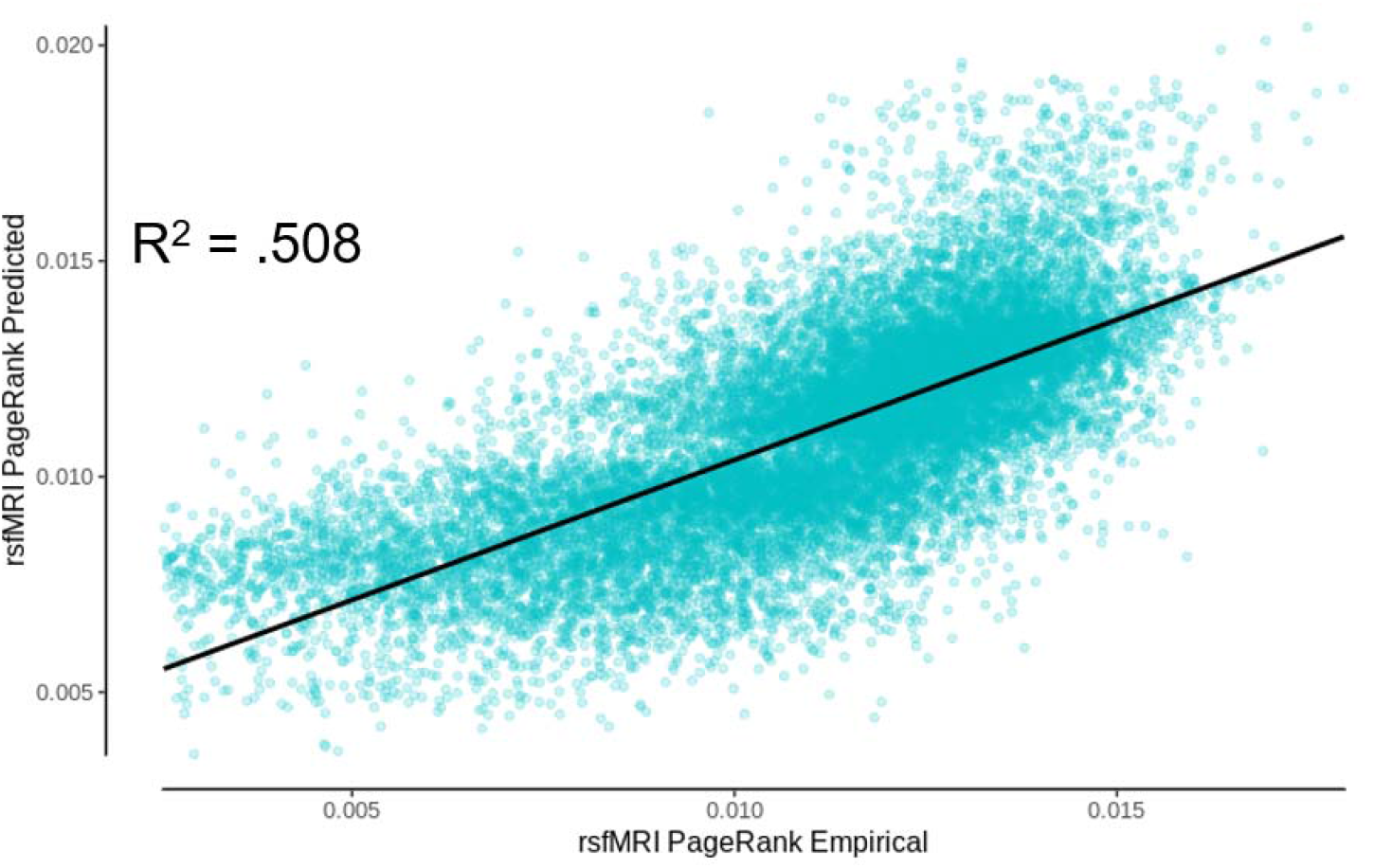
AAL atlas non-aggregated rsfMRI functional PageRank centrality measures calculated from the predicted functional connectivity values as a function of empirical rsfMRI functional PageRank centrality (R^2^ = .508).

**Supplementary Figure 9.**
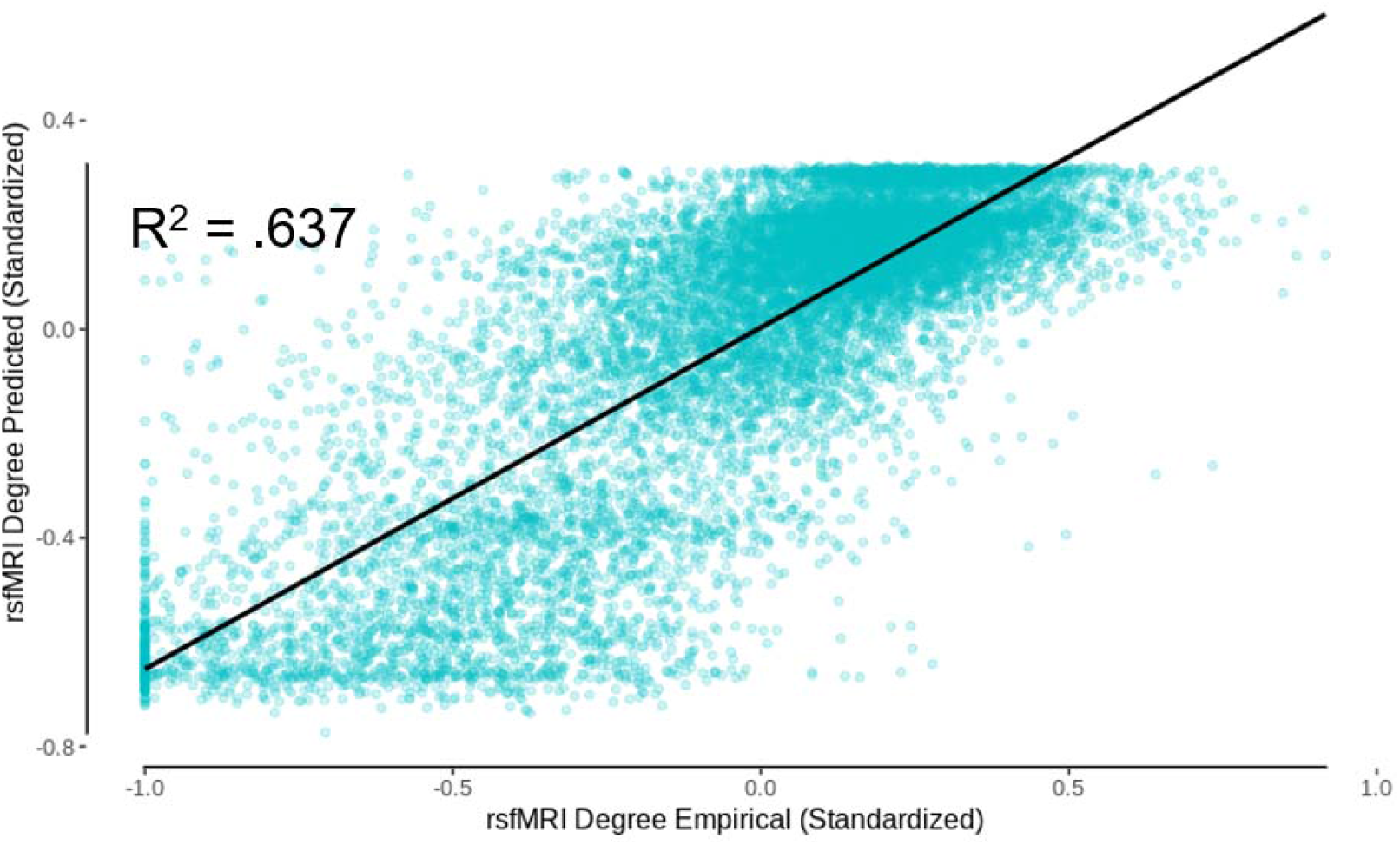
DK atlas non-aggregated predicted rsfMRI functional connectivity degree centrality as a function of empirical rsfMRI functional degree centrality (R^2^ = .637).

**Supplementary Figure 10.**
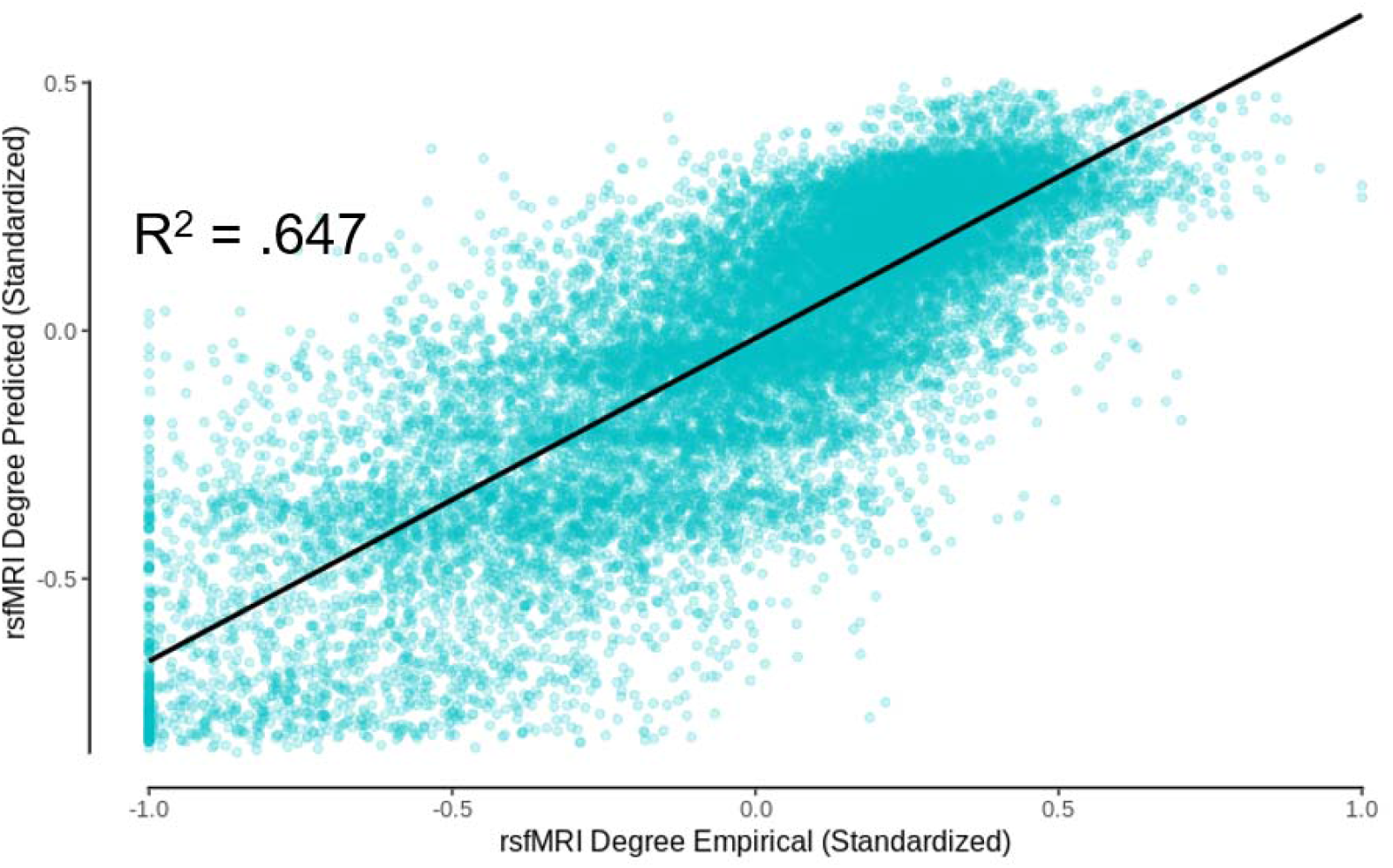
AAL atlas non-aggregated predicted rsfMRI functional connectivity degree centrality as a function of empirical rsfMRI functional degree centrality (R^2^ = .647).

**Supplementary Figure 11.**
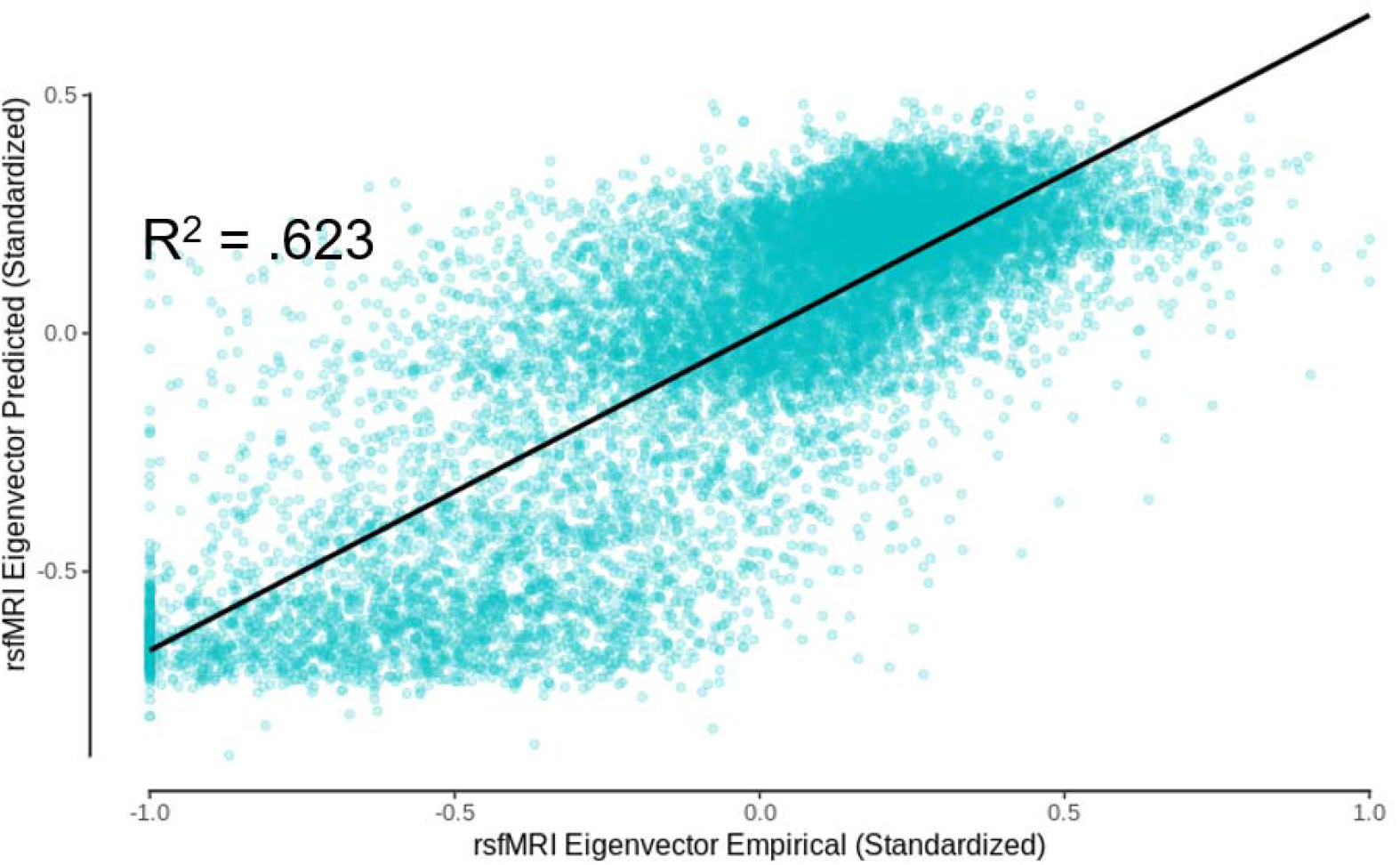
DK atlas non-aggregated predicted rsfMRI functional connectivity eigenvector centrality as a function of empirical rsfMRI functional eigenvector centrality (R^2^ = .623).

**Supplementary Figure 12.**
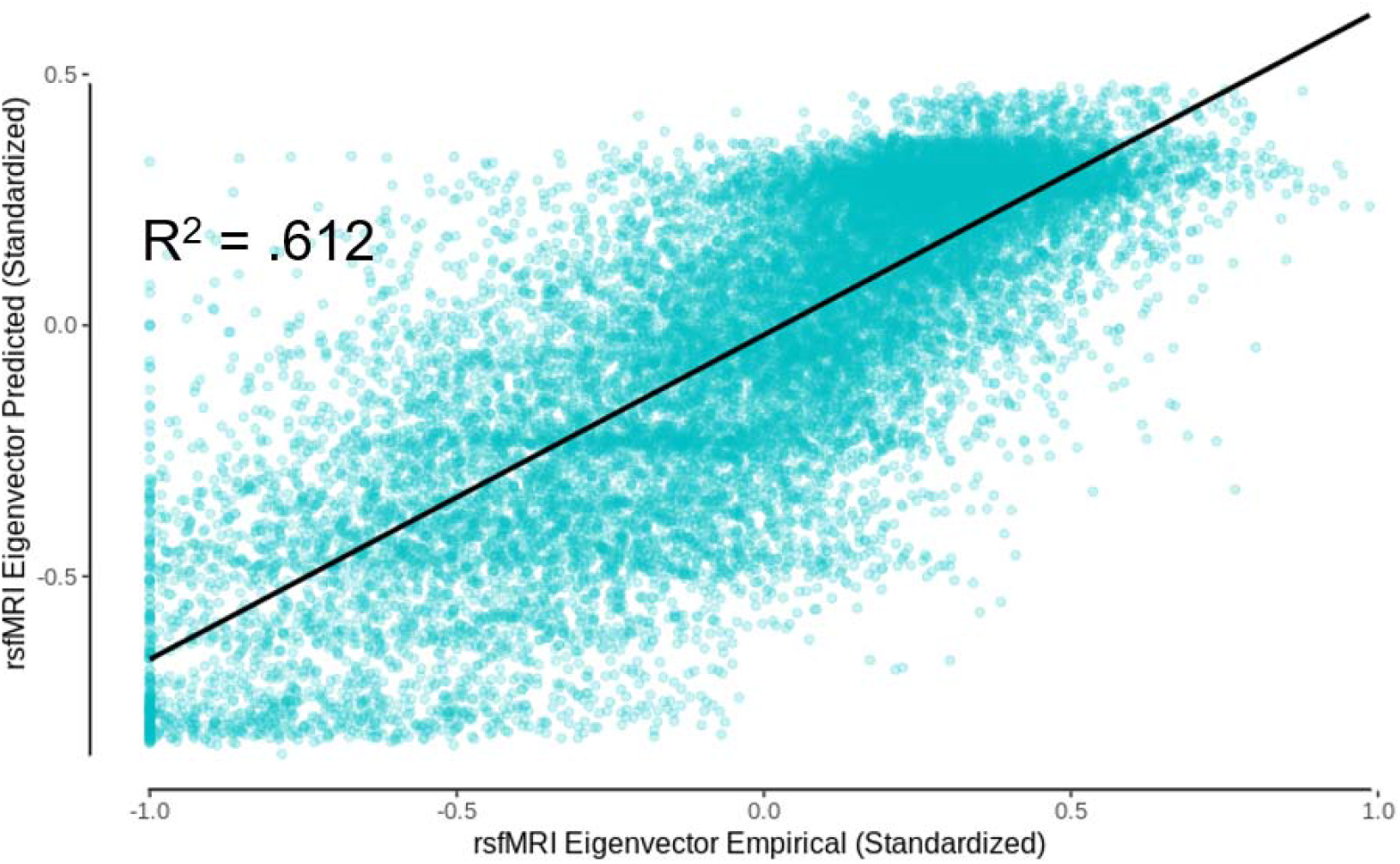
AAL atlas non-aggregated predicted rsfMRI functional connectivity eigenvector centrality as a function of empirical rsfMRI functional eigenvector centrality (R^2^ = .612).

**Supplementary Figure 13.**
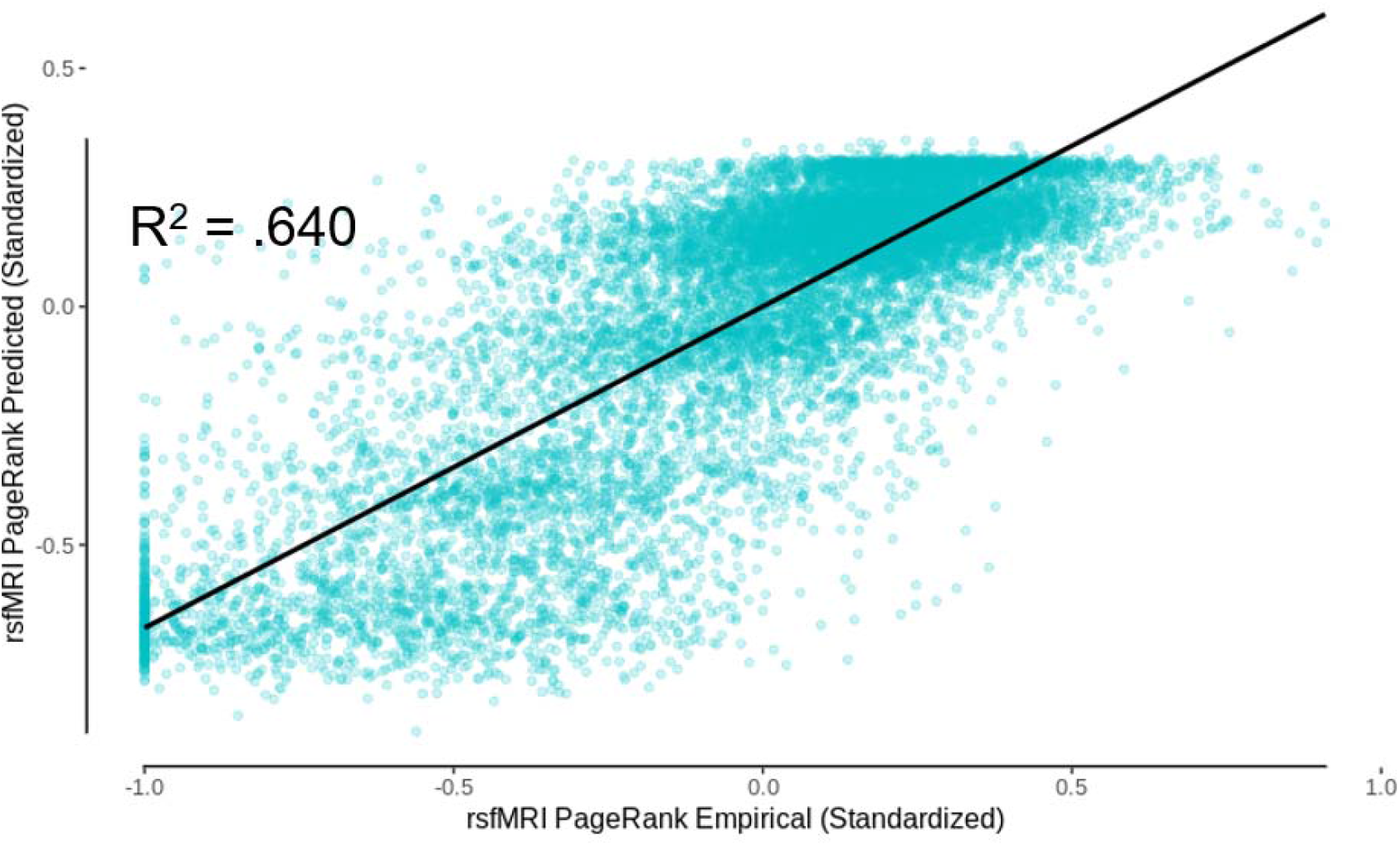
DK atlas non-aggregated predicted rsfMRI functional connectivity PageRank centrality as a function of empirical rsfMRI functional PageRank centrality (R^2^ = .640).

**Supplementary Figure 14.**
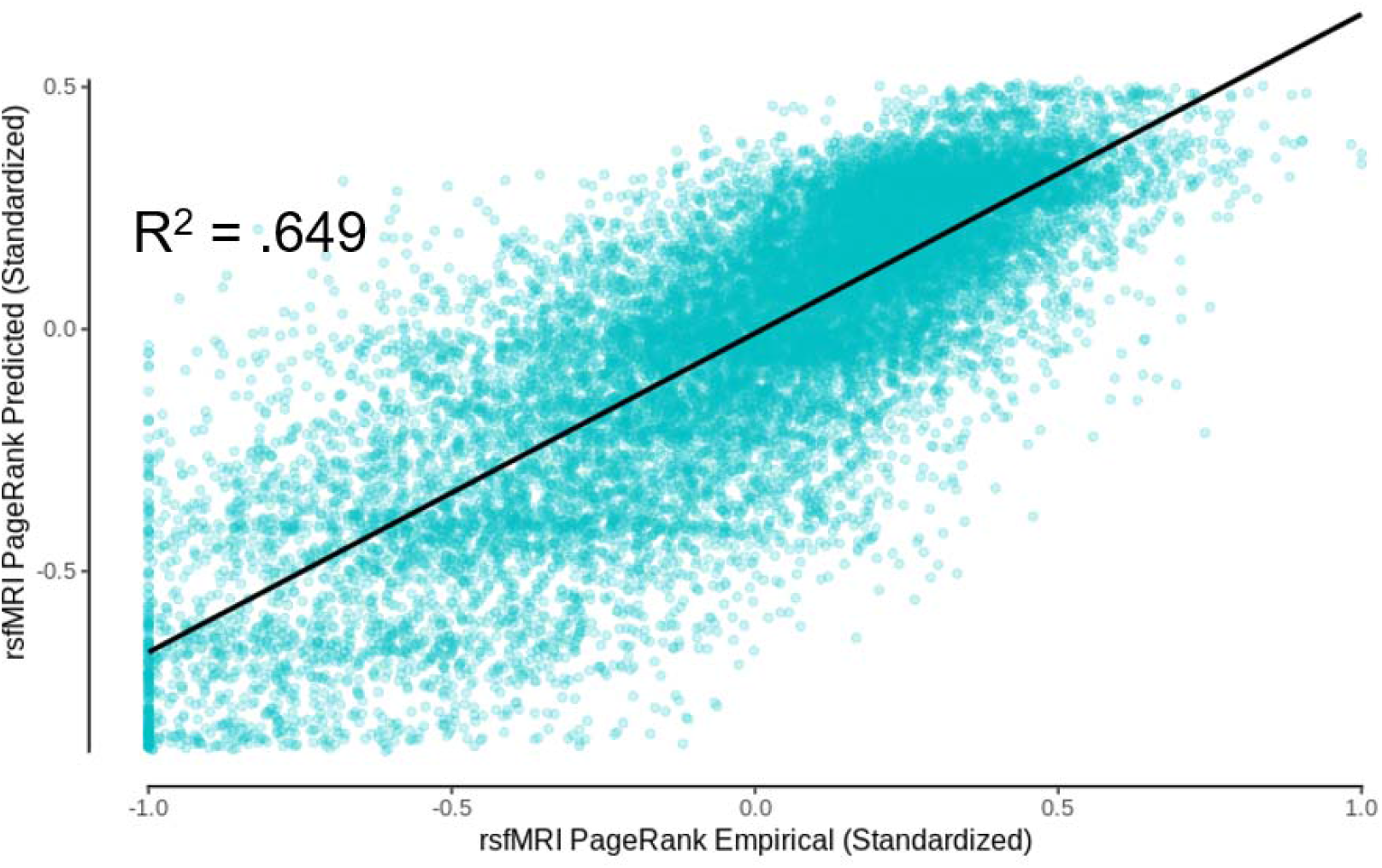
AAL atlas non-aggregated predicted rsfMRI functional connectivity PageRank centrality as a function of empirical rsfMRI functional PageRank centrality (R^2^ = .649).

